# COCO-Search18: A Dataset for Predicting Goal-directed Attention Control

**DOI:** 10.1101/2020.07.27.221499

**Authors:** Yupei Chen, Zhibo Yang, Seoyoung Ahn, Dimitris Samaras, Minh Hoai, Gregory Zelinsky

## Abstract

Attention control is a basic behavioral process that has been studied for decades. The currently best models of attention control are deep networks trained on free-viewing behavior to predict bottom-up attention control—saliency. We introduce COCO-Search18, the first dataset of laboratory-quality *goal-directed behavior* large enough to train deep-network models. We collected eye-movement behavior from 10 people searching for each of 18 target-object categories in 6202 natural-scene images, yielding *∼*300,000 search fixations. We thoroughly characterize COCO-Search18, and benchmark it using three machine-learning methods: a ResNet50 object detector, a ResNet50 trained on fixation-density maps, and an inverse-reinforcement-learning model trained on behavioral search scanpaths. Models were also trained/tested on images transformed to approximate a foveated retina, a fundamental biological constraint. These models, each having a different reliance on behavioral training, collectively comprise the new state-of-the-art in predicting goal-directed search fixations. Our expectation is that future work using COCO-Search18 will far surpass these initial efforts, finding applications in domains ranging from human-computer interactive systems that can anticipate a person’s intent and render assistance to the potentially early identification of attention-related clinical disorders (ADHD, PTSD, phobia) based on deviation from neurotypical fixation behavior.

---

The control of visual attention comes broadly in two forms. One is bottom-up, where control is exerted purely by the visual input^1, 2^. This is the form of attention predicted by saliency models, which exploded in popularity in the behavioral fixation-prediction and computer-vision literatures^1, 3 –5^. The other form of control is top-down, where behavioral goals rather than bottom-up salience control the allocation of visual attention. Goal-directed attention control underlies all the things that we *try* to do, and this diversity makes its prediction vastly more challenging than predicting bottom-up saliency, and more important. In addition to its basic research value, a better understanding of goal-directed attention could lead to the development of biomarkers for neurotypical attention behavior against which clinical conditions can be quantitatively compared, and to advances in intelligent human-computer interactive systems that can anticipate a user’s visual goals and render real-time assistance^6 –8^.

Goal-directed attention has been studied for decades^9 –16^, largely in the context of visual search. Search is arguably the ost basic of goal-directed behaviors; there is a target object and the goal is to find it, or conclude its absence. Goals are extremely effective in controlling the allocation of gaze. Imagine two encounters with a kitchen, first with the goal of learning the time from a wall clock and again with the goal of warming a cup of coffee. These “clock” and “microwave” searches would yield two very different patterns of eye movement, as recently demonstrated in a test of this gedanken experiment^17^, and understanding this goal-directed control has been a core aim of search theory. The visual search literature is itself voluminous (see reviews^18–20^). Here we focus on the prediction of image locations that people fixate as they search for objects, and how the selection of these fixation locations depends on the target goal.

The visual search literature is not only mature in its empirical work, it is also rich with many hugely influential theories and models^12–16, 21^. Yet despite this success, over the last years progress has stalled. Our premise is that this is due to the absence of a dataset of search behavior sufficiently large to train deep network models. Our belief is based on observation of what occurred in the bottom-up attention-control literature during the same time. The prediction of fixations during free viewing, the task-less cousin of visual search, has become an extremely active research topic, complete with managed competitions and leaderboards for the most predictive models^22^ (http://saliency.mit.edu/). The best of these saliency models are all deep networks, and to our point, all of them were trained on large datasets of labeled human behavior^23–27^. For example, one of the best of these models, DeepGaze II^23^, is a deep network pre-trained on SALICON^25^. SALICON is a crowd-sourced dataset consisting of images that were annotated with mouse-based data approximating the attention shifts made during free viewing. This model of fixation prediction during free viewing was therefore trained on a form of free-viewing behavior. Without SALICON, DeepGaze II, and models like it^24–27^, would not have been possible, and our understanding of free-viewing behavior, widely believed to reflect bottom-up attention control, would be greatly diminished. For the task of visual search, there is nothing remotely comparable to SALICON^25^. Here we describe in detail COCO-Search18, the largest dataset of goal-directed search fixations in the world. COCO-Search18 was recently introduced at CVPR2020^28^, and our aim in this paper is to elaborate on the richness of this dataset so as to increase its usefulness to researchers interesting in modeling top-down attention control.

## Methods

### Behavioral Data Collection

COCO-Search18 is built from Microsoft COCO, Common Objects in Context^29^. COCO consists of over 200,000 images of scenes that have been hand-segmented into 80 object categories. This ground-truth labeling of objects in images makes COCO valuable for training computer vision models of object detection^29 –33^. However, in order for COCO to be similarly valuable for training models of goal-directed attention, these images would also need to be labeled with the locations fixated by people searching for different target-object goals. COCO-Search18 fills this niche by providing these training labels of search behavior.

The dataset consists of a large-scale annotation of a subset of COCO, 18 of its 80 object categories, with goal-directed search fixations. Each of 10 participants searched for each of 18 target-object categories (blocked) in 6,202 COCO imges, mostly of indoor scenes. This effort required an average of 12 hours per participant, distributed over 6 days. This substantial behavioral commitment makes it possible to train models of individual searchers^28^, although our focus here is on group behavior. The eye position of each participant was sampled every millisecond using a high-quality eye-tracker under controlled laboratory conditions and procedure, resulting in *∼*70,000,000 gaze-position samples in total. These raw gaze samples were clustered into 299,037 search fixations (*∼*30,000 per participant), which dropped to 268,760 fixations after excluding those from incorrect trials. Figure 1 shows representative images and fixation behavior for each target category. See SM1 for details about: selection criteria (for images, target categories, and fixations), the eye tracker and eye tracking procedure, participant instruction, and a comparison between COCO-Search18 and existing datasets of search behavior.

**Figure 1.**
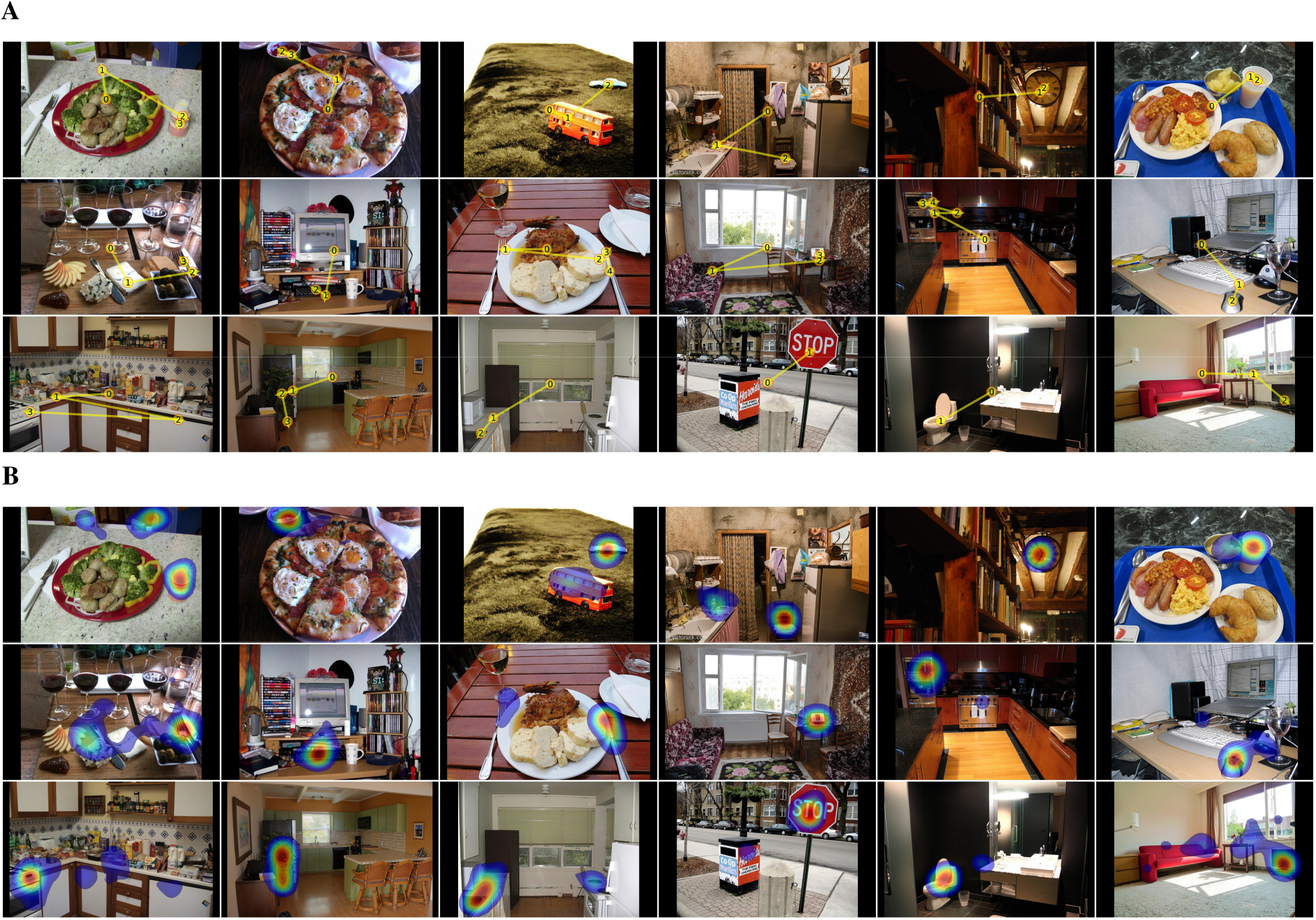
(A). Examples of target-present images for each of the 18 target categories. Yellow lines and numbered discs indicate a representative search scanpath from a single participant. From left to right, top to bottom: bottle, bowl, car, chair, (analog) clock, cup, fork, keyboard, knife, laptop, microwave, mouse, oven, potted plant, sink, stop sign, toilet, tv. (B). Examples of fixation density maps (excluding initial fixations at the center) computed over participants for the same scenes.

### Search-Relevant Image Statistics

Figure 2A shows three search-relevant characterizations of the COCO-Search18 images. The left panel shows the distribution of target-object sizes, based on bounding-box COCO labels. This distribution skewed toward smaller targets, with the range constrained by image selection to be between 1% and 10% of the image size (see SM1). The mean visual angle of the targets, based on the square root of bounding-box size, was 8.4°, about the size of a clenched fist at arm’s length. The middle panel shows the distribution of initial target eccentricities, which is how far the target appeared in peripheral vision, based on center fixation at the start of search. Target eccentricities ranged from 10° to 25° of visual angle, with a mean of *∼*15° eccentricity. The right panel shows the distribution of the number of “things” in each image, again based on the COCO object and stuff labels^34^. Some images depicted only a handful of objects, whereas others depicted 20 or more (keeping in mind that this labeling was coarse). We report this statistic because search efficiency is known to degrade with the number of items in a search array^35^, and a similar relationship has been suggested for the feature and object clutter of scenes^36–38^. Figure 2B again shows these measures, now grouped by the 18 target categories. Target size and initial target eccentricity varied little across target categories, while the measure of set size varied more. See SM2 for analyses showing how each of these three measures correlated with search efficiency, for each target category.

**Figure 2.**
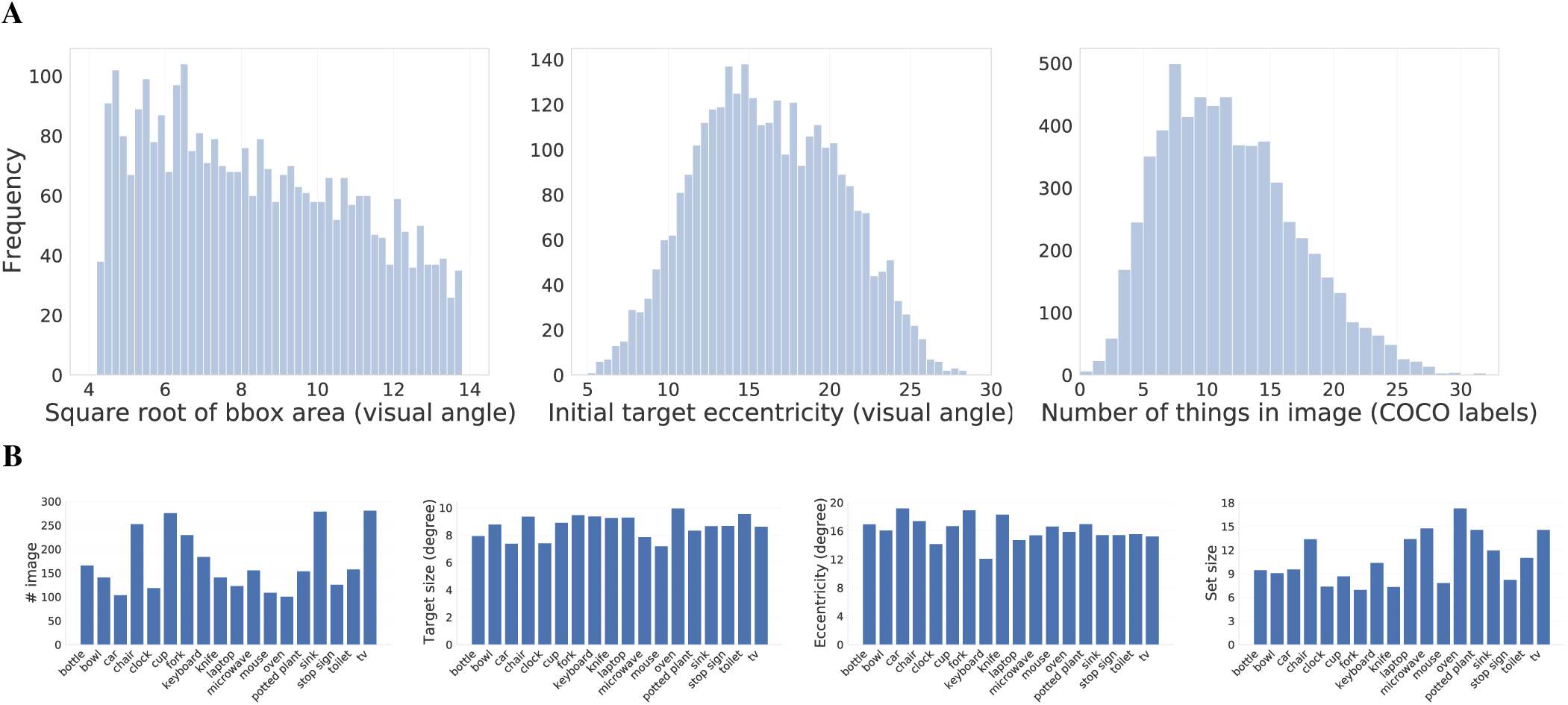
(A). Distributions of target sizes, based on the visual angle of their bounding-box areas (left), and initial target eccentricities (middle), both for the target-present images. The number of “things” (objects and “stuff” categories, both based on COCO-stuff labels) appearing in the search images (right). (B). Image statistics from COCO-Search18, grouped by the 18 target categories. The left plot shows the number of images, followed by three analyses paralleling those presented in (A): averaged target-object size in degrees of visual angle, initial target eccentricity based on bounding-box centers, and the average number of things in an image (a proxy for set size).

### Search Procedure and Metrics

The paradigm used for data collection was speeded categorical search^39–41^. The participant’s task was to indicate whether an exemplar of a target category appeared in an image of a scene (Figure S3). They did this by making a target present/ absent judgment as quickly as possible while maintaining accuracy. The target category was designated at the start of a block of trials. Half of the search images depicted an exemplar of a target (target-present, TP), and the other half did not (target-absent, TA).

We measure goal-directed attention control as the efficiency in which gaze moves to the search target. Because the target was an object category, the term used for this measure of search efficiency is *categorical target guidance*^39,41^, defined as the controlled direction of gaze to a target-category goal. We consider multiple measures of target guidance in Figure 3, but here we focus on the cumulative probability of fixating the target after each search saccade^42–45^. A target category that can successfully control gaze will be fixated in fewer eye movements compared to one that has less capacity for target guidance. A desirable property of the target-fixation-probability (TFP) function (Figure 4) is that it is meaningful to compute the area under the TFP curve (TFP-auc), which we suggest as a new metric for evaluating search guidance across target categories and models.

**Figure 3.**
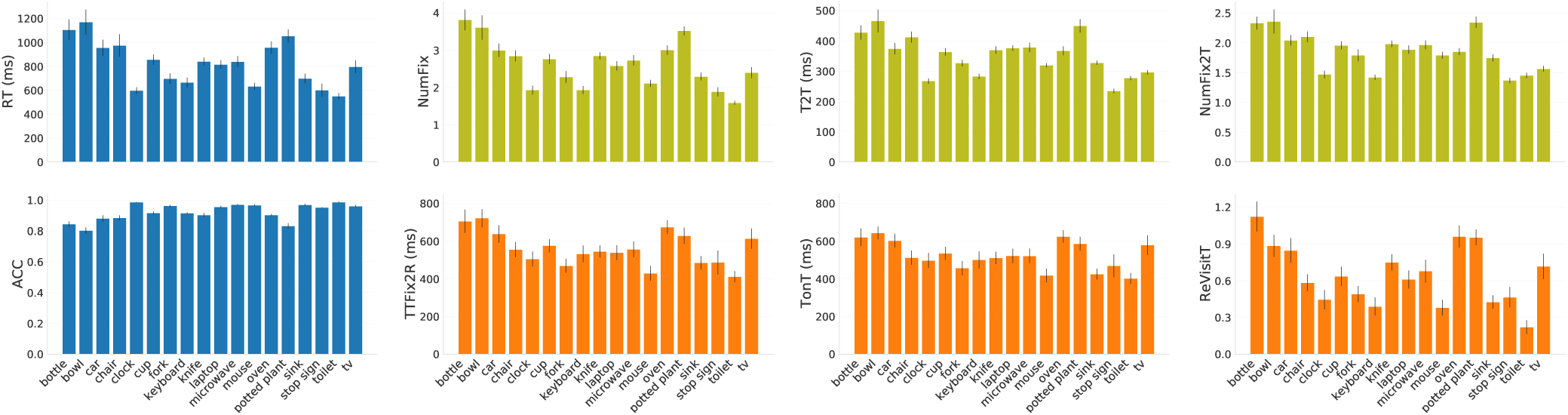
Basic behavioral analyses of the target-present data from COCO-Search18, grouped by the 18 target categories. Blue plots (left two) show the manual measures of reaction time (RT) and response accuracy (ACC). Olive plots (top row) show gaze-based analyses of categorical guidance efficiency: number of fixations made before the button press (NumFix), time until first target fixation (T2T), and number of fixations made until first target fixation (NumFix2T). Orange plots (bottom row) show gaze-based measures of target verification: time from first target fixation until response (TTFix2R), total time spent fixating the target (TonT), and the number of re-fixations on the target (ReVisitT). Values are means over 10 participants, and error bars represent standard errors.

**Figure 4.**
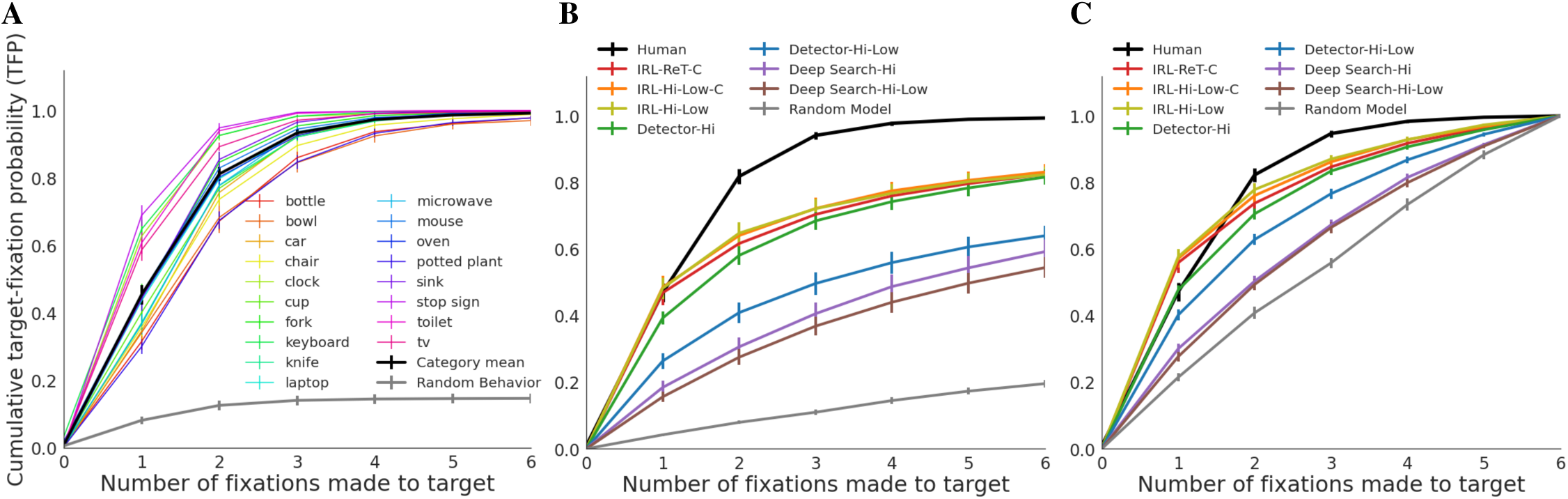
(A). Cumulative probability of fixating the target (y-axis; target-fixation probability or TFP) as a function of fixation serial position (x-axis; 0-6), shown individually for the 18 target categories (color lines) and averaged over target types (bold black line). The bottom-most function is a Random Behavior baseline obtained by computing target-fixation probability using a scanpath from the same participant searching for the same target category but in a different image. For the 18 target functions, means were computed by first averaging over images and then over participants, and standard errors were computed over participants. For the averaged behavioral data and the Random Behavior baseline (black and gray lines), means were computed by first averaging over images and then over categories, and standard errors were computed over categories. (B). TFP functions generated from model predictions on the test images. Names designate a model type (IRL, Detector, Deep Search) and a state representation (ReT, Hi-Low, Hi, C), separated by hyphens. Average behavioral TFP is again plotted in bold black, this time for just the test data (which explains the small differences from the corresponding function in A, which included the training and testing data). The Random Model baseline was obtained by making six movements of the Hi-Low foveated retina, with ISTs after each, and determining whether any of these movements brought the high-resolution central window to the target. Means were first computed over images and then over categories, and standard errors were computed over categories. (C). A re-plot of B, but only including data from trials in which the target was successfully fixated within the first six fixations (i.e., search scanpaths that succeed in locating the target.)

### Model Comparison

Now that COCO-Search18 exists, what can we do with it? To answer this question we conducted benchmarking to deter-mine how well current state-of-the-art methods, using COCO-Search18, can predict categorical search fixations. To create a context for this model comparison we considered three very different modeling approaches, which all shared a common backbone model architecture, a ResNet50 pre-trained on ImageNet^46^.

Our first approach predicted search fixations using object detectors trained for each of the target categories. We did this by re-training the pre-trained ResNet50 on just the 18 target categories using the COCO labels. Standard data augmentation methods of re-sizing and random crops were used to increase variability in the training samples. We then used these trained detectors to predict search fixations on the test images. For a given target and test image, we obtained a confidence map from the target detector and used it to sample a sequence of fixation locations based on the level of confidence. Note that this approach is pure computer vision, meaning that it uses the image pixels solely and knows nothing about behavior.

With COCO-Search18, however, it is possible to also train on the search behavior. There are multiple ways of doing this. In our second approach we re-trained the same ResNet50, only this time using labels as the fixations made by searchers viewing the training images. Specifically, fixation-density maps (FDMs) were obtained for each TP training image for a given category, and these were used as labels for model training. This model is in a sense a search version of models like DeepGaze II^23^ in the free-viewing fixation-prediction literature, which are also trained to predict FDMs. We therefore refer to this model as Deep Search. Deep Search differs from the Target Detector model in that it is trained on search fixation density to predict search behavior.

For our third modeling approach we used inverse-reinforcement learning (IRL)^47–49^, an imitation-learning method from the machine-learning literature, to simply mimic the search scanpaths observed during training. We chose IRL over other imitation-learning methods because it is based on reward, known to be a powerful driver of behavior^50–52^, but we think it is likely that other imitation-learning methods would perform similarly. The IRL model we used^49^ works by learning, through an adversarial process playing out over many iterations, how to make model-generated behavior, initially random, become more like human behavior. It does this by rewarding behavior that happens to be more human-like. IRL is therefore very different from a Target Detector, but also different from Deep Search, which also gets to use search behavior in its training. The IRL model learns to imitate the search scanpath, meaning the sequence of fixations made to the search target, whereas Deep Search uses only FDMs that do not represent the temporal order of fixations. Because the IRL model used the most search behavior for training, we hypothesized that it would best predict search behavior in our model comparison. See SM3 for additional details about IRL.

### State Comparison

In addition to the model comparison, we also compared several state representations used by the models. In the current context, the state is the information that is available to control search behavior, and essential to this are the features extracted from each search image. We refer to the original images as high-resolution (Hi-Res), in reference to the fact that they were not blurred to reflect retina constraints. Extracting features from a Hi-Res image produces a Hi-Res state, and it is this state that is used by most object-detection models in computer vision where the goal is to maximize detection success. Primate vision, however, is profoundly degraded from this Hi-Res state by virtue of the fact that we have a foveated retina. A foveated retina means that high-resolution visual inputs exist only for a small patch of the image at the current fixation location, and blurred everywhere else. Given our goal to model the fixation behavior of the COCO-Search18 searchers, each of whom had a foveated retina, we included this basic biological constraint in the state to determine its importance in model training and prediction of search behavior (see also^53^). Relatedly, and as fundamentally, each new fixation changes the state by allowing high-resolution information to be obtained from the vantage of a new image location. Capturing these fixation-dependent spatio-temporal state changes in the context of search was a core goal in the development of COCO-Search18.

We considered two fovea-inspired states. In the first we used the method from Perry and Geisler^54^ to compute a Retina-Transformed (ReT) image. A ReT image is a version of the Hi-Res image that is blurred to approximate the gradual loss in visual acuity that occurs when viewing at increasing eccentricities in peripheral vision. Second, we implemented an even more simplified foveated retina consisting of just a high-resolution central patch (7° × 7° visual angle) surrounded by low-resolution “peripheral” vision elsewhere, with the critical difference from the ReT image being that only a single level of blur (Gaussian filter with *σ* = 2) was used to approximate the low-resolution periphery. Computing the gradual blur used in the ReT image was computationally very demanding, and the inclusion of the simpler Hi-Low state was motivated largely to reduce these computational demands (ReT requires *∼*15× the processing time per image). However, having this condition lso enabled a needed initial evaluation of how veridically low-level visual-system constraints need to be followed when training deep-network models of human goal-directed behavior.

We also considered two spatio-temporal state representations for how information is accumulated with each new fixation in a search scanpath. A behavioral consequence of having a foveated retina is that we make saccadic eye movements, and the order in which these eye movements are made correspond to different visual states. Our first spatio-temporal state assumed a high-resolution foveal window that simply moves within a blurred image. This means that each change in fixation brings peripherally blurred visual inputs into clearer view, and causes previously clear visual inputs to become blurred. This spatio-temporal state representation is aligned most closely with the neuroanatomy of the oculomotor system, so we will consider this to be the default state. However, this default state representation assumes that foveally-obtained information on fixation *n* is completely lost by fixation *n+1*, and indeed something like this is true for high-resolution information about visual detail^55^. However, this state fails to capture any memory for the fixated objects that persists over eye movement, which is also known to exist^56^. To address the potential for an object context to build over fixations, we therefore also used a state that accumulates the high-resolution foveal views obtained at each fixation in the search scanpath, a state we refer to as Cumulative (-C). Over the course of multi-fixation search, the Hi-Low-C state would therefore accumulate high-resolution foveal snapshots with each new fixation, progressively de-blurring what would be an initially moderately-blurred version of the image. We explore these two extremes of information preservation during search so as to inform future uses of a fovea-inspired spatio-temporal state representation to train deep network models.

### General Model Methods

All of the models followed the same general pipeline. Each 1050 × 1680 image input was resized to 320 × 512 pixels to reduce computation. This is what we refer to as the Hi-Res image (or just Hi); the ReT and Hi-Low images were computed from this. These images were passed through the ResNet50 backbone to obtain 20 × 32 feature map outputs, with the features extracted from these images now reflecting either Hi-Res, ReT, or Hi-Low states, respectively. Different models were trained using these features and others, as described in the Model Comparison section, and all model evaluations were based on a 70% training, 10% validation, and 20% testing, random split of COCO-Search18 within each target category. See SM3 for additional details about the training and testing separation, and Figure S15 for how the two compare on search performance measures.

The trained models were used to obtain model-specific priority maps for the purpose of predicting the search fixations in each test image. The priority map for the Target Detector was a map of detector confidence values at each pixel location, and fixations were sampled probabilistically from this confidence map.The priority map for Deep Search is its prediction of the FDM, given the input image and the model’s learned mapping between image features and the FDM ground-truth during training. The priority map for the IRL model is the reward map recovered during its training, which recall occurred during its learning to mimic search behavior. Because this search behavior was itself reward driven, the priority map for the IRL model is therefore a map of the total reward expected by making a sequence of search fixations to different locations in a test image. The IRL model was additionally constrained to have an action space discretized into a 20 × 32 grid, which again was done to reduce computation time. A given action, here a change in fixation, is therefore a selection of 1 from 640 possible grid cells, a sort of limitation imposed on the spatial resolution of the model’s oculomotor system. The selected cell was then mapped back into 320 × 512 image space by upscaling, and the center of this cell became the location of the model’s next fixation. The non-IRL models made their action selection directly in the 320 × 512 image space, with higher priority values selected with higher probability.

All of the model × state combinations in our comparison were required to make six changes in fixation for each test image. This number was informed by the behavioral data showing that the probability of target fixation was clearly at ceiling by the sixth eye movement (Figure 4A). To produce these 6-fixation scanpaths, we iterated the fixation generation procedure using inhibitory spatial tagging (IST), which is a mechanism serving the dual functions of (1) breaking current fixation, thereby enabling gaze to move elsewhere, and (2) discouraging the refixation of previously searched locations. IST has long been used by computational models of free viewing^57,58^ and search^59,60^. Here we enforce IST by setting the priority map to zero after each fixation over a region having a radius of 2.5° visual angle (based on a 3 × 3 grid within the 20 × 32 action space). IST was applied identically after each fixation made by all of the models. This was true even for models that did not have a foveated retina, such as a Target Detector with a Hi-Res state, in which case IST was applied to the image locations selected for “fixation”. See SM3 for additional details.

The nomenclature that we adopted for the model comparison consists of the model type as the base and the state representation as a suffix. If the spatio-temporal state is cumulative, there is a second suffix of -C. For example, the IRL-ReT-C model accumulates graded-resolution foveal views of an image with each reward-driven eye movement. Although our aim is to explore as systematically as possible each state for every model, for some models a given state representation is not applicable. For example, it makes no sense for the IRL model to use the Hi-Res state. Because that state representation does not change from one search fixation to the next it would be impossible to learn fixation-dependent changes in state, thereby defeating the purpose of using the IRL method. Similarly, it makes no sense to have a cumulative state for anything but the IRL model, as the others would be unable to use this information. However, it does make sense to test a Target Detector and Deep Search on a Hi-Low state as well as a Hi-Res state, and these models are included in the Table 1 model evaluation.

**Table 1.** Results from fixation-prediction models (rows) using multiple scanpath metrics (columns) applied to the COCO-Search18 test images. Arrows indicate the direction of better prediction success, and values in bold indicate best predictions across the model comparison. In the case of Sequence Score and MultiMatch, “Human” refers to an oracle method whereby one searcher’s scanpath is used to predict another searcher’s scanpath; “Human” for all other metrics refers to observed behavior. See the main text for additional details about the scanpath-comparison metrics, and SM3 for purely spatial comparisons using the AUC, NSS, and CC metrics.

| | TFP-AUC ↑ | Probability<br>Mismatch ↓ | Scanpath<br>Ratio ↑ | Sequence<br>Score ↑ | MultiMatch ↑ | | | |
| --- | --- | --- | --- | --- | --- | --- | --- | --- |
|  |  |  |  |  | shape | direction | length | position |
| Human | 5.200 | - | 0.862 | 0.489 | 0.903 | 0.736 | 0.880 | 0.910 |
| Random Model | 0.744 | 4.455 | 0.392 | 0.266 | 0.832 | 0.579 | 0.783 | 0.755 |
| Detector-Hi | 4.001 | 1.209 | 0.680 | 0.411 | 0.877 | 0.665 | 0.837 | 0.872 |
| Detector-Hi-Low | 2.975 | 2.225 | 0.601 | 0.370 | 0.863 | 0.640 | 0.820 | 0.833 |
| Deep Search-Hi | 2.519 | 2.681 | 0.579 | 0.348 | **0.890** | 0.627 | **0.867** | 0.861 |
| Deep Search-Hi-Low | 2.282 | 2.918 | 0.546 | 0.333 | 0.882 | 0.617 | 0.859 | 0.848 |
| IRL-ReT-C | 4.170 | 1.131 | 0.731 | 0.418 | 0.879 | 0.673 | 0.842 | 0.874 |
| IRL-Hi-Low-C | **4.262** | **1.031** | 0.747 | **0.419** | 0.886 | **0.677** | 0.849 | **0.885** |
| IRL-Hi-Low | 4.245 | 1.036 | **0.753** | 0.417 | 0.884 | **0.677** | 0.847 | **0.885** |

## Results

### Behavioral Performance

We interrogated COCO-Search18 using multiple performance measures. Figure 3 reports these analyses for each of the target categories. Analyses can be conceptually grouped into manual measures (accuracy and response time; blue plots), gaze-based measures of categorical guidance (number of fixations before the button press, and both the time and number of fixations until the first target fixation; olive plots), and measures of target verification time (time from first target fixation until the button press, total time spent fixating the target, and the number of target re-fixations; orange plots). What is clear from these analyses is that, except for accuracy, there is wide variability across target categories in these measures, and this variability creates fertile ground for future model development. Also clear from Figure 3 is that there is considerable correlation among some of these measures, perhaps most evident among the search guidance measures where the shapes of the plots look similar. We include these different measures, not to suggest their independence, but rather as a courtesy to readers who may be familiar with different measures.

Figure 5 is a matrix visualization of these analyses, now with color coding a ranking of search efficiency. In Figure 5A, the deepest red for each measure (row) indicates the least efficient (or most difficult) search over the 18 target categories, and the deepest blue indicates the most efficient (or easiest) search. The appearance of columns in this visualization captures the agreement among the measures. More subtle patterns in the data can also be seen. For example, the two predominately red columns at the left indicate agreement in that the bottle and bowl objects were difficult targets, speculatively because these target categories have particularly high variability in their image exemplars. Relatedly, appearing near the right are two of the consistently easiest targets, stop signs and toilets, both having relatively well-defined category membership. Figure 5B shows a similar plot, only now performance is averaged over target categories and plotted for individual participants. Search accuracy and efficiency clearly differ among the participants in this ranking. Participants 7 and 8 were better searchers than Participants 2 and 9, meaning that they tended to find the target faster and with fewer fixations while keeping a low error rate. Differences in search strategy can also be seen from this visualization. Participant 1 searched the display carefully, resulting in few missed targets, but this person’s search was not very efficient. In contrast, Participant 4 was quick to find and verify targets, but had relatively low accuracy. See SM2 for parallel analyses of the target-absent data from COCO-Search18.

**Figure 5.**
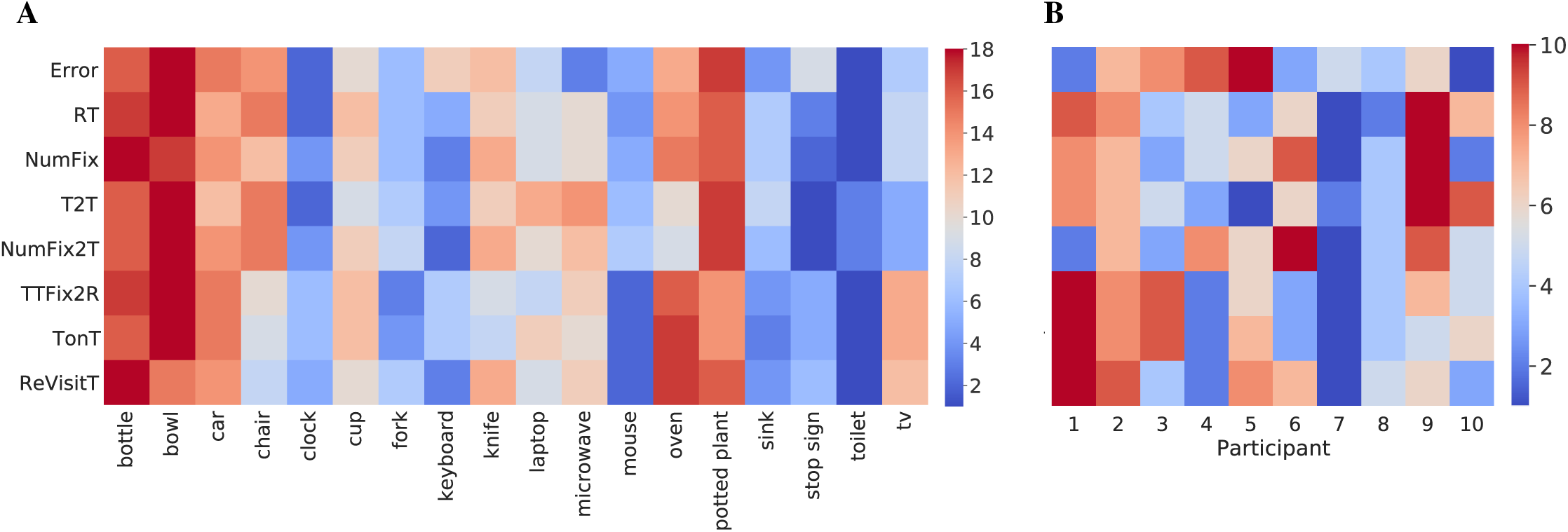
(A). Ranked target-category search efficiency [1-18], averaging over participants. Redder color indicates higher rank and harder search targets, bluer color indicates lower rank and easier search. Target category is grouped (columns) and shown for multiple performance measures (rows). These measures include: response Error, reaction time (RT), number of fixations (NumFix), time to target (T2T), number of fixations to target (NumFix2T), time from first target fixation until response (TTFix2R), time spent fixating the target (TonT), and the number of target re-fixations (ReVisitT). (B). A similar ranking of the target-present data, only now for participant efficiency (columns 1-10), averaged over target category. Performance measures and color coding are the same as in panel A.

However, arguably the gold-standard measure of attention control is the cumulative probability of fixating the target after each saccade made during search, the target-fixation probability (TFP). Figure 4A shows TFP functions for the first six search saccades, averaged over participants and plotted for individual target categories. The mean behavior over targets is indicated by the bold black function. The noteworthy pattern is that the slope of the group function is far steeper than that of the chance baseline, obtained by computing TFP using a scanpath from a different image but from the same participant and target category. On average, about half of the targets were fixated with the very first saccade. By the second saccade TFP jumped to .82, and by the third saccade it reached a performance ceiling of .94, which increased only slightly after saccades 4–6. This high degree of attention control means that, although we aimed to create a search dataset having a moderate level of difficulty, COCO-Search18 skews easy. This is due in part to unexpectedly large practice effects (see SM2 for details). However, it is fortuitous that such a strong attention-control signal exists in the behavioral data, given the challenge faced by even start-of-the-art models in predicting this simple search behavior.

### Model Evaluation

Fixation prediction models broadly fall into two groups, models that predict the spatial distribution of locations fixated by participants viewing an image (i.e., the FDM), and models that predict both the location and order of the fixations made by a person viewing an image (i.e., the scanpaths). In a search task, fixation behavior changes dramatically over the first few eye movements ^61^, making it important to consider the spatio-temporal fixation order. For this reason, we will focus on spatio-temporal fixation prediction here and defer discussion of purely spatial FDM prediction to SM4, and especially Table S1. Both types of prediction were based on the 6-saccade sequences that each model was required to make for each test image. Specifically, 10 6-fixation scanpaths (excluding the initial fixation) were predicted for each test image by sampling probabilistically from the generated priority map, and for each of these search scanpaths the model behavior was analyzed up to first fixation on the target, or six changes in fixation, whichever came first.

Predicting the spatio-temporal order of search fixations can also take two forms. One has been to make fixation predictions with respect to the search target. For example, predicting the probability of the target being fixated by the first search saccade, the second, etc. These target-based predictions capture the efficiency of search, where the goal is to find the target, and models making this type of prediction have been the more common in the search literature ^45,62^. Here we use three metrics to evaluate the success of these predictions. Two of these metrics were derived from the TFP function (Figure 4A): TFP-auc, which is the area under the cumulative target-fixation-probability curve, and Probability Mismatch, which sums over each fixation in a scanpath the absolute differences between the behavioral and model TFP. The third metric, Scanpath Ratio, is the Euclidean distance between the initial fixation location (roughly the center of the image) and the location of the target (center of bounding box) divided by the summed Euclidean distances between the fixation locations in the search scanpath^42^. It is a search efficiency metric because an initial saccade that lands directly on the target would yield a Scanpath Ratio of 1, and all less efficient searches would be *<* 1. An alternative to predicting target guidance over the spatio-temporal search scanpath is to predict the scanpath itself. This approach assumes that any target guidance would be reflected in the sequence of fixated image locations leading up to the target decision. We considered two metrics for comparing behavioral and predicted search scanpaths: Sequence Score, which clusters scanpaths into strings and uses a string matching algorithm for comparison^63^, and MultiMatch, which takes a multi-dimensional approach to computing scanpath similarity^64,65^. Both metrics capture properties of the spatio-temporal search scanpath and place less importance on the fact that there is a search target. SM4 should be consulted for additional details about these metrics.

Table 1 provides an evaluation of how each model×state combination fared in fair comparison using these metrics. As we hypothesized, the three IRL models generally outperformed the others (see Table S2 for statistical tests). They did so for every metric except MultiMatch, where all the models performed similarly. The only other model that was comparably predictive was Detector-Hi, but this model has no fovea and is therefore the least biologically plausible. A perhaps clearer picture of this model comparison can be obtained by comparing the behavioral TFP function to ones computed for each model. Figure 4B shows this evaluation of search efficiency for each of the model × state combinations (in color) and for the mean search behavior (in black), limited to the TP test data. Focusing first on state comparisons, we did not find large differences between the states tested. Whether blur was graded or binary appeared not to matter, as indicated by the very similar TFP functions for the ReT and the Hi-Low states using the IRL model. This pattern also appeared in Table 1, where the IRL models differed by tiny margins. For this reason, and its far greater computational efficiency, we adopted only the Hi-Low state in the other model comparisons (therefore, there are no Deep Search-ReT or Detector-ReT models). Similarly, but specific to the IRL model, it made little difference whether or not the state accumulated high-resolution visual information with each fixation in a search scanpath. The fact that the IRL model seemed not to use this accumulated visual information is broadly consistent with the view that very little high-resolution information is preserved across saccades^55^. However, it did matter whether the state included a foveated retina or not, as exemplified by the difference between Hi-Res and Hi-Low states for the Detector model. This state comparison suggests that future work may want to avoid manipulations of fine-grained retinal blur and assumptions about intersaccadic visual memory, and focus on adding more basic limitations on human visual perception to a model’s pipeline, with the inclusion of a Hi-Low foveated retina being one example.

All of the tested models made reasonable predictions of search behavior in this challenging benchmark, where “reasonable” is liberally defined as bearing greater resemblance to the human behavior than the chance baseline. However, the Deep Search models and the Detector-Hi-Low model were clearly less efficient in their search behavior than either human behavior or any of the IRL models. This poor relative performance is likely caused by these models not capturing the serial order of search fixations, and that this order maters. A corollary finding is that the IRL models, because they learned these spatio-temporal sequences of search fixations, better predicted search behavior. This was true for all the IRL models, which all predicted the efficiency of the first search fixation almost perfectly (IRL models vs. Human at fixation 1 with post-hoc t-tests, all *ps*_*bonferroni*_ = 1.0). Also interesting is the degree that an object detector (Detector-Hi) can predict search behavior, supporting previous speculation^66^. If an application’s goal is to predict a person’s early fixation behavior during search without regard for biological plausibility, a simple object detector will work well based on our testing with COCO-Search18. Another finding from Figure 4B is that none of the models achieved the high level of successful target fixation exhibited in human performance. Performance ceilings after six saccades (termed *fixated-in-6 accuracy*) ranged from .54 (Deep Search-Hi-Low) to .83 (IRL-Hi-Low-C), all well below the near perfect fixated-in-6 accuracy (.99) from human searchers (post-hoc t-tests with all *ps*_*bonferroni*_ <.001). These lower performance plateaus, undoubtedly reflecting limitations in current object detection methods, means that the models tended either to fixate the target efficiently in the first one or two eye movements (like people), or tended not to fixate the target at all (unlike people). If a model cannot represent the features used for target guidance as robustly as people, there may be images for which there is essentially no guidance signal, and on these inefficient search trials the number of eye movements needed to fixate the target would often be greater than six, hence the performance plateaus.

These different performance ceilings are problematic in that they conflate limitations arising from object detection with limitations in effective target prioritization, as measured by search efficiency. For example, a strength of the TFP-auc metric is that it is grounded in the TFP functions from Figure 4B, but this means that it includes the different performance ceilings in its measure and this weakens it as a pure measure of attention control. To address this concern, in Figure 4C we again plot TFP functions, but now only for trials in which the target was successfully fixated within the first six saccades. By restricting analysis to only trials having perfect fixated-in-6 accuracy, the metric becomes more focused on search efficiency. By this measure, and keeping in mind that the data are now skewed toward easier searches, the IRL-Hi-Low-C and IRL-Hi-Low models remain the most predictive overall, although now all IRL models overestimate slightly the efficiently of the first search saccade. But perhaps the biggest winner in this comparison is the Detector-Hi model, which now predicts TFP almost perfectly after the first fixation, and has generally improved performance for subsequent fixations. We tentatively conclude that simple prioritization of fixations by an object detector predicts reasonably well the prioritization of behavioral fixations in visual search. The losers in this comparison were the Deep Search models, which remained less efficient than human behavior even after normalization for fixated-in-6 accuracy.

## Discussion

Recent years taught us the importance of large datasets for model prediction, and this importance extends to models of attention control. COCO-Search18 is currently the largest dataset of goal-directed search fixations, having sufficient number to be used as labels for training deep network models. We conducted a systematic (but still incomplete) exploration of models and state representations to provide some initial context for the types of model predictions that are possible using COCO-Search18, given current state-of-the-art (or nearly so). This model comparison focused on the degree that search behavior was used during training, ranging from none (Detector), to some (Deep Search), to entire search-fixation scanpaths (IRL). With respect to the IRL model, its use with COCO-Search18 is the first attempt to predict the spatio-temporal movements of goal-directed attention by training on human search behavior. We found that the IRL model was far more predictive of search efficiency than the Detector-Hi-Low model or either of the Deep Search models, despite the Deep Search models using methods considered to be state-of-the-art in the fixation-prediction literature on free-viewing behavior. In our state comparison we focused on the different ways that a primate foveated retina, and its movement, might be represented and used to train fixation prediction models. We also extensively benchmarked COCO-Search18, both in terms of the search behavior that it elicited, analyzed using multiple behavioral measures and metrics, and in terms of the predictive success of models ranging in their degree of training on the COCO-Search18 behavior. All this means that COCO-Search18 can be used immediately to start generating new testable hypotheses. But likely the greatest contribution of this work is yet to come. With a dataset the size and quality of COCO-Search18, opportunities exist to explore new policies and reward functions for predicting goal-directed control that have never before been possible ^28^. Our hope is that COCO-Search18 will strengthen the bridge that human attention has built between the machine learning and behavioral science literatures.

COCO-Search18 is now part of the MIT/Tuebingen Saliency Benchmark, previously the MIT Saliency Benchmark but renamed to reflect the group that is now managing the competition. The training, validation, and test images in COCO-Search18 are already freely available as part of COCO^29^. Researchers are also free to see and use COCO-Search18’s training and validation search fixations, but the fixations on the test images are withheld. As part of a managed benchmark, in a separate track it will be possible to upload predictions and have them evaluated on this test dataset. We invite you to participate in this good-natured adversarial competition, and we hope that you enjoy using COCO-Search18: https://github.com/cvlab-stonybrook/Scanpath_Prediction.

## Supplementary Materials

### SM1: Behavioral Data Collection

#### Comparable datasets of search behavior

Figure S1 shows how COCO-Search18 compares to other large-scale datasets of search behavior. To our knowledge, there were only three such image datasets that were annotated with human search fixations^17,67,68^. In terms of number of fixations, number of target categories, and number of images, COCO-Search18 is far larger. The PET dataset^68^ collected search fixations for six animal target categories in 4,135 images selected from the Pascal VOC 2012 dataset^69^, but the search task was non-standard in that participants were asked to “find all the animals” rather than search for a particular target category. This paradigm is therefore search at the super-ordinate categorical level, which is far more weakly guided than basic-level search^70^. Gaze fixations were also recorded for only 2 seconds/image, and multiple targets often appeared in each scene. The microwave-clock search dataset (MCS^17^) is our own work and a predecessor of COCO-Search18. In collecting data for the 18 target categories in COCO-Search18 we had to start somewhere, and our first two categories were microwaves and clocks (although the datasets differed for even those two categories due to the use of different exclusion criteria). Until recently, perhaps the best dataset of search fixations was from^67^, but it is relatively small, limited to only the search for people in scenes, and is now a decade old. Note that, whereas there are larger datasets with respect to free-viewing fixations (SALICON^25^) or fixations collected using other visual tasks (POET^71^), these tasks were not visual search and therefore these datasets cannot be used to train models of search behavior. These collective inadequacies demanded the creation of a newer, larger, and higher-quality dataset of search fixations, enabling deep network models to be trained on people’s movements of attention as they pursue target-object goals.

#### Selection of target categories and search images

Here we more fully describe how we selected from COCO’s trainval dataset^29^ the 18 target categories and the 6,202 images included in COCO-Search18. A goal in implementing our selection criteria was to elicit the behavior that we are trying to measure, namely, the guidance of search fixations by a target category. We also put care into excluding images that might elicit other gaze patterns that would introduce noise with respect to identifying the target-control signal. This sort of attention to detail is uncommon in datasets created for the training of deep network models, where the approach seems to be “the more images the better”. But whereas this is usually true because more images leads to better-trained models, in creating a dataset of human behavior this more-is-better impulse should be tempered with some quality control to be confident that the behavior is of the purported type. In the current context this behavior should be search fixations that are guided to the target, because search fixations that are un-guided have less value as training labels. Because a standard search paradigm collects behavioral responses for both TP and TA images, separate selection criteria were needed. All image selection was based on object labels and/or bounding boxes provided by COCO. On this point, while inspecting the images that were ultimately selected we noticed that exemplars in some categories were mislabeled, probably due to poor rater agreement on that category. For instance, several chair exemplars were mislabeled as couches, and vice versa. Rather than attempting to correct these mislabels, which would be altering COCO, we decided to keep them and tolerate a higher-than-normal error rate for the affected categories. This action seemed best, given our plan to discard error trials from the search performance analyses in our study, but researchers interested in interpreting button press errors in COCO-Search18 should be aware of this labeling issue.

##### Target-present image selection

Six criteria were imposed on the selection of images to be used for target-present search trials.

1. Images were excluded if they depicted people or animals. We did this to avoid the known biases to fixate on these objects when they appear in a scene^22,72^. Such biases would compete with guidance from target-category features, thereby distorting study of the target-bias that is more central to search.
2. Images were excluded if they depicted multiple instances of the target. A scene showing a classroom with many chairs would therefore be excluded from the “chair” target category because one, and only one, instance of a chair would be allowed in an image.
3. Images were excluded if the size of the target, measured by the area of its bounding box, was smaller than 1% or larger than 10% of the total image area. This was done to create searches that were not too hard or too easy.
4. Images were excluded if the target appeared at the image center, based on a 5 × 5 grid. We did this because the participant’s gaze was pre-positioned at this central location at the start of each search trial.
5. Images were excluded if their width/height ratio fell outside the range of 1.2-2.0 (based on a screen ratio of 1.6). This criterion excluded very elongated images, which we thought might distort normal viewing behavior.
6. Images, and entire image categories, were excluded if the above criteria left fewer than 100 images per object category. We did this because fewer than 100 images would likely be insufficient for training and testing a deep network model specific to that object category.

Applying these exclusion criteria left 32 object categories from COCO’s original 80. Given that this left still far too many images for people to practically annotate with search fixations, we decided to attempt exclusion of images where targets were highly occluded or otherwise difficult to recognize. We did this out of concern that such images would largely introduce noise into the search behavior. To do this, we trained object detectors on cropped views of these 32 categories, and excluded images if the object bounding boxes had a classification confidence < .99. Specifically, for these 32 categories we created a validation set consisting of images meeting the selection criteria and a training set consisting of the images that did not. The bounding box of the object, for each of the 32 object classes, was then cropped in the image to obtain the positive training samples. Negative samples were same-sized image patches that had 25% intersection with the target (area of intersection divided by area of target), meaning that they were class-specific hard negatives. All cropped patches (over 1 million) were resized to 224 × 224 pixels while maintaining the aspect ratio using padding. The classifier was a ResNet50 pre-trained on ImageNet, which we fine-tuned by dilating the last fully-connected layer and re-training on 33 outputs (32+”Negative”). Images were excluded if the cropped object patch had a classification score of less than .99. This procedure resulted in 18 categories with at least 100 images in each category, totaling 3,131 TP images.

Two final exclusion criteria were implemented by manual selection. First, for the clock target category we included only images of analog clocks, meaning that we excluded digital clocks from being clock targets. We did this because the features of analog and digital clocks are highly distinct and very different, and we were concerned that this would introduce variability in the search behavior and reduce data quality. Five images depicting only digital clocks were excluded for this reason. Lastly, images from all 18 of the target categories were screened for objectionable content, which we defined as offensive content or content evoking discomfort or disgust. The “toilet” category had the most images (17) excluded for objectionable content, with a total of 25 images excluded across all target categories. After implementing all exclusion criteria discussed in this section, we obtained 3,101 TP images from 18 categories: bottle, bowl, car, chair, (analog) clock, cup, fork, keyboard, knife, laptop, microwave, (computer) mouse, oven, potted plant, sink, stop sign, toilet, and tv. See Figure 2 for the specific number of images in each category.

##### Target-absent image selection

To balance the selection of the 3,101 TP images, we selected an equal number of TA images from COCO. To do this, we kept the criteria excluding images depicting people or animals, extreme width/height image ratios, and images with objectionable content, all as described for the TP image selection, but added two more exclusion criteria that were specific to each of the 18 target-object categories.

1. Images were excluded if they depicted an instance of the target, a prerequisite for a TA image.
2. Images were excluded if they depicted less than two instances of the target category’s siblings, a criterion introduced to discourage searchers from making TA responses purely on the basis of scene type. For example, a person might be biased to make a TA response if they are searching for a toilet target and the image is a street scene. Because COCO has a hierarchical organization, parent, child, and sibling relationships can be used for image selection. For example, COCO defines the siblings of a microwave to be an oven, toaster, refrigerator, and sink, all under the parent category of appliance. By requiring that the TA scenes for a target category have at least two of that category’s siblings, we impose a sort of scene constraint that minimizes target-scene inconsistency and makes a scene appropriate to use as a TA image. A scene that has an oven and a refrigerator is very likely to be a kitchen, thereby making it difficult to answer on the basis of scene type alone whether a microwave target is present or absent.

These exclusion criteria still left us with many thousands more TA images than we needed, so we sampled randomly within each of the 18 target categories to match the 3,101 TP images.

##### Order of target-category presentation

Collecting the search behavior for 6,202 images required dividing each participant’s effort into six days of testing. Each testing session was conducted on a different day, lasted about 2 hours, and consisted of about 1000 search trials, evenly divided between TP and TA. Because images from different categories can overlap (e.g., images depicting a microwave may also depict an oven), the presentation order of the target-category blocks was constrained to minimize the repetition of images in consecutive categories and consecutive sessions. For example, because 49 images satisfied the selection criteria for both the sink and microwave target categories, we prevented the microwave and sink categories from appearing in, not only the same session, but the sessions preceding and following. We did this to minimize possible biases resulting from seeing the same scene in different search contexts. A heuristic for maximizing this distance between repeating images resulted in the following fixed target category presentation order across the six sessions:

1. tv + sink;
2. fork + chair;
3. car + bowl + potted plant + mouse;
4. knife + keyboard + oven + clock;
5. cup + laptop + toilet;
6. bottle + stop sign + microwave.

Each participant viewed from Session 1 to Session 6, or from Session 6 to Session 1, with this order counterbalanced across participants.

#### Data-collection procedure

Participants were 10 Stony Brook University undergraduate and graduate students, 6 males and 4 females, with ages ranging from 18–30 years. All had normal or corrected to normal vision, by self report, were naive with respect to task design and paradigm when recruited, and were compensated with course credit or money for their participation. Informed consent was obtained from each participant at the beginning of testing, in accordance with the Institutional Review Board responsible for overseeing human-subjects research at Stony Brook University.

The target category was designated to participants at the start of each block. This was done using the type of display shown in Figure S2 for the potted-plant and analog clock categories. The name of the target category was shown in text at the top, with examples of objects that would, or would not, qualify as exemplars of the named category. In selecting exemplars to illustrate as positive target-category members, we attempted to capture key categorical distinctions at a level immediately subordinate to the target category. When needed, we also gave negative examples by placing a red X through the object. We did this to minimize potential confusions and to enable the participant to better define the target category’s boundary.

The procedure (Figure S3) on each trial began with a fixation dot appearing at the center of the screen. To start a trial, the participant would press the “X” button on a game-pad controller while carefully looking at the fixation dot. An image of a scene would then be displayed and the participant’s task would be to answer, “yes” or “no”, whether an exemplar of the target category appears in the displayed scene by pressing the right or left triggers of the game-pad, respectively. The search scene remained visible until the manual response. Participants were told that there were an equal number of TP and TA trials, and that they should make their responses as fast as possible while maintaining high accuracy. No accuracy or response time feedback was provided.

The presentation of images during the experiment was controlled by Experiment Builder (SR research Ltd., Ottawa, Ontario, Canada). Stimuli were presented to participants on a 22-inch LCD monitor (1680 × 1050 pixel resolution) at a viewing distance of 47cm from the monitor, enforced by chin and head rests. These viewing conditions resulting in horizontal and vertical visual angles of 54°× 35°, respectively. Participants were asked to keep their gaze on the fixation point at the start of each trial, but were told that they should feel free to move their eyes as they searched. Eye movements were recorded throughout the experiment using an EyeLink 1000 eye-tracker in tower-mount configuration (SR research Ltd., Ottawa, Ontario, Canada). Eye-tracker calibrations occurred before every block or whenever necessary, and these 9-point calibrations were not accepted unless the average calibration error was *≤*.51° and the maximal error was *≤*.94°. The experiment was conducted in a quiet laboratory room under dim lighting conditions.

### SM2: Behavioral evaluation of COCO-Search18

#### Effects of set size and target eccentricity

The visual search literature has done excellent work in identifying many of the factors that increase search difficulty (for reviews, see:^12,18,60,73^). Larger set sizes (number of items in the search display), smaller target size, larger target eccentricity, and greater target-distractor similarity are all known to make search more difficult. However, most of this work was done in the context of simple stimuli, and generalization to realistic images is challenging. For example, what to consider an object in a scene is often unclear, making it difficult to define a set size^74^. Objects in images also do not usually come annotated with labels and bounding boxes. These problems of object segmentation and identification, which largely do not exist for search studies using object arrays, become significant obstacles to research when scaled up to images of scenes.

With COCO-Search18, we can begin to ask how the search for targets in images is affected by set size and target eccentricity. Set size is determined based on the COCO object and stuff labels, which collectively map every pixel in an image to an object or stuff category. Set size is the count of the number of these labels for a given image. Figure S4 shows the relationship between the number of fixations made on an image, averaged over participants, and the set size of that image, grouped by target category. Some target categories, such as laptop, oven, microwave, and potted-plant, have significant positive set size effects (*r* = .21 to .37, *p*s *≤*.01), indicating a less efficient search with more objects. A similar pattern is shown in Figure S5 for the relationship between the number of fixations on a search image and the initial visual eccentricity of the target (distance between the image center and the target bounding-box center), where for these same objects there was a decrease in search efficiency with increasing target eccentricity. For other target object categories, such as: stop sign, fork, and keyboard, search efficiency was unaffected by either set size or target eccentricity (*p*s *>* .05), possibly because these objects are either highly salient (stop sign) or highly constrained by scene context (keyboard).

#### Distance between search fixations and the target

How much closer does each search fixation bring gaze to the target? We analyzed this measure of search efficiency and report the results in Figure S6. Plotted is the Euclidean distance between the target location and the locations of the starting fixation (0) and the fixation locations after the first six eye movements (1-6). The most salient pattern is the rapid decrease in fixation-target distance in the first two new fixations, which dovetails perfectly with the steep increase in the cumulative probability of target fixation over these same eye movements reported in Figure 4A. From a starting location near the center of the image, these eye movements brought gaze steadily closer to the target. Note that because this fixation-target distance is averaged over images and participants, the roughly 5 degrees of visual angle at the bottom of these functions should not be misinterpreted as gaze being this distance from the target on a given trial. More interpretable are the overall trends, where a steep drop in distance is followed by a plateau, or even a smaller increase in distance with the 5th and 6th new fixations. This small increase is likely an artifact of these 5 and 6-fixation trials being the most difficult, with more idiosyncratic search behavior.

#### Target-absent search fixations

In the main text we focused on the TP data, where the guidance signal is clearer and the modeling goals are better defined, but we conducted largely parallel analyses of the TA data. Figure S7A shows representative TA images with fixation data from one participant, and Figure S7B shows FDMs from all participants for the same images. Comparing these data with the TP data from Figure 1, it is clear that people made many more fixations in the absence of a target. This was expected from the search literature, but it should also be noted that the FDMs are still much sparser than what would be hypothesized by an exhaustive search. Paralleling Figure 3, in Figure S8 we report applicable analyses of the TA search behavior. These are grouped by manual accuracy and response time, and the mean number of fixations made before the target-absent button press terminating a trial. Note that accuracy was high (low false positive error rate) for all of the target categories except chairs and cups, with the reason for the former already discussed in the context of mislabeling and the reason for the latter likely reflecting an occasionally challenging category distinction (e.g., some bottles can look like some cups). Also note that there was an average of only five fixations made during search, even on the TA search trials. As in Figure 5, Figure S9 visualizes the agreement and other patterns among these measures. The rows show ranked performance, with dark red indicating more difficult (or least efficient) search and dark blue indicating relatively easy or efficient search. The columns in Figure S9A group the measures by target category. Similar to the TP data, there was again good consistency among the measures. Also consistent is the fact that bottles and cups were among the most difficult target categories, whereas the toilet category was the easiest. There was also evidence in the TA data for a speed-accuracy trade-off for some target categories. For example, microwaves and stop signs had relatively low error rates, but these categories were searched with relatively high effort, as measured by ranked response time and number of fixations. Figure S9B visualizes the measures by participant instead of category, where we again found individual differences between participants in search efficiency.

#### Practice effects

Each of the participants contributing to COCO-Search18 searched more than 6000 images, making it possible to analyze how their search efficiency improved with practice. Figure S10 shows practice effects for both response time (top) and the number of fixations before the button press (bottom), where we define practice effects as performance on the first 1/3 of the trials compared to performance on the last 1/3 of the trials for each target category. Practice effects were larger for TA trials (right) than for TP trials (left), noting the differences in y-axes scales, and that considerable differences existed across categories. Some categories, such as bottles, showed large practice effects, while other categories, such as analog clocks, showed none at all. We speculate that this difference is due to some categories requiring more exemplars to fully learn compared to others. For example, analog clock was perhaps the most well defined of COCO-Search18’s categories, and bottle certainly one of the least well defined, creating greater opportunity to better learn the bottle category with practice over trials.

#### Search fixation durations

Figures S11 and S12 show density histograms of the search fixation durations for the TP and TA data, respectively, plotted for each of the target categories. Fixation durations are plotted across the x-axes with a bin size of 50ms, and y-axes show the normalized probability density at each fixation. Of note in the TP data is that the mode initial fixation durations (blue lines) were a bit longer than the mode duration of the rest (averaged mode difference = 63ms), consistent with the very strong guidance observed in the initial eye movements, and they tended to have more bi-modal distributions. The main peak was at *∼*250 ms, with a smaller and very short-latency peak at *∼*50 ms that is likely a truncation artifact of fixation duration being measured relative to the onset of the search display. In contrast, the distributions of second fixations (orange lines) were consistently shorter, even relative to the subsequent fixations. Speculatively, this may be due to a greater proportion of the first new fixations being “off object”^75^, which are often followed by short-latency corrective saccades that bring gaze accurately to an object. This interpretation is consistent with the high probability of the target being fixated by the second eye movement (Figure 4A). As for the subsequent fixations, they tended to be short (*∼*200ms) and not highly variable in their durations. The TA fixations showed similar trends, except for the durations of the second fixations no longer differing from the rest.

#### Saccade amplitudes

We also analyzed the distribution of saccade amplitudes during visual search, defined here as the Euclidean distance between consecutive fixations in visual angle. Figure S13 and Figure S14 show the distributions of saccade amplitudes in the TP and TA data, respectively. In the TP data, saccade amplitudes were larger in some categories (toilet and stop sign) than others (bottle and potted plant), likely because easier target categories could be identified from farther in the visual periphery. There was also evidence for bimodality in the amplitude distributions, shown most clearly for clocks, forks, stop signs, and tvs. We speculate that this bimodality reflects larger-amplitude exploratory saccades mixed with smaller-amplitude saccades used in the verification of an object category. Mean saccade amplitudes in the TA data were clearly larger than for the TP data (*t*(17) = 11.79, *p <* .001), and this difference was consistent across target categories (all *p*s *≤*.001). We attribute this to the relatively large viewing angle of the search displays (54 × 35 degrees of visual angle) creating a greater need for exploration, but this is also speculation. The distributions of saccade amplitudes were also more consistent across categories in the TA data, with there being weaker evidence of bi-modality.

### SM3: Model Methods

#### Training and testing datasets

Model success depends on the training dataset being an accurate reflection of the test dataset. When the training dataset includes a behavioral annotation, as does COCO-Search18, it is therefore important to know that similar patterns exist in the training and testing search behavior. The analyses shown in Figure 5A included images from all of COCO-Search18, which recall were randomly split into 70% for training, 10% for validation, and 20% for testing. Figure S15 replots the data from Figure 5A, but divides it into the training/validation (left) and testing (right) datasets. Note the high agreement between the testing and train/val datasets across this battery of behavioral performance measures.

#### Inverse Reinforcement Learning

The specific inverse-reinforcement learning (IRL) method that we used was generative adversarial imitation learning (GAIL^49^) with proximal policy optimization (PPO)^76^. The model policy is a generator that aims to create state-action pairs that are similar to human behavior. The reward function (the logarithm of the discriminator output) maps a state-action pair to a numeric value. The generator and discriminator are trained within an adversarial optimization framework to obtain the policy and reward functions. The discriminator’s task is to distinguish whether a state-action pair was generated by a person (real) or by the generator (fake), with the generator aiming to fool the discriminator by maximizing the similarity between its state-action pairs and those from people. The reward function and policy that are learned from the fixation-annotated images during training are then used to predict new search fixations in the unseen test images.

### SM4: Performance metrics and model evaluation

#### Metrics for comparing search efficiency and scanpaths

We considered five metrics for quantifying search efficiency and comparing search scanpaths (Table 1). Two metrics for quantifying search efficiency follow directly from the group target-fixation probability (TFP) function shown in Figure 4. The first of these computes the area under the TFP curve, a metric we refer to as TFP-auc. Second, and relatedly, we compute the sum of the absolute differences between the human and model target-fixation-probabilities in a metric that we refer to as Probability Mismatch. A third metric for quantifying overt search efficiency is Scanpath Ratio. It is the Euclidean distance between the initial fixation location and the target divided by the summed Euclidean distances between the fixation locations in the search scanpath^42^. It is an efficiency metric because an initial saccade that lands directly on the target would give a Scanpath Ratio of 1, meaning that the distance between starting fixation and the target would be the same as the summed saccade distance. These three metrics emphasize target-fixation efficiency by penalizing either the number of fixations or the saccade-distance traveled to achieve the target goal. The final two metrics focus on scanpath comparison, and specifically comparing the search scanpaths between people and the models. The first of these scanpath-comparison metrics computes a Sequence Score by first converting a scanpath into a string of fixation cluster IDs, and then using a string matching algorithm^63^ to measure the similarity between the two strings. Figure S16 shows examples of behavioral and model scanpaths and their sequence scores to develop an intuition for this metric. Lastly, we use MultiMatch^64,65^ to measure the scanpath similarity at the pixel level. MultiMatch measures five aspects of scanpath similarity: shape, direction, length, position, and duration. We excluded the duration measure from our use of this metric because the models in our comparison group did not predict fixation duration. See Table S2 for the results of statistical tests comparing predictions from each pair of models.

#### Comparing predicted and behavioral fixation-density maps (FDMs)

Search has a temporal dynamic, making a metric for capturing the spatio-temporal sequence of fixations preferred over ones that compare only FDMs, where this temporal component is disregarded. However, the prediction of FDMs is common for free-viewing tasks, and because there is no technical reason why FDM metrics cannot be applied to search we do so here in the hope that the visual saliency literature finds this comparison useful. Models generated scanpaths having a maximum length of 6 new fixations, but FDMs were constructed only from those fixations leading up to the first fixation on the target, just as FDMs were constructed from the behavioral fixations. We used three widely accepted metrics for comparing predicted against observed FDMs. Area Under the Receiver Operating Characteristic Curve (AUC) uses a predicted priority map as a binary classifier to discriminate behavioral fixation locations from non-fixated locations. Normalized Scanpath Saliency (NSS) finds the model predictions at each of the behavioral fixation locations, then averages and normalizes these values. Lastly we computed a Pearson’s Correlation Coefficient (CC) between the predicted and behavioral FDMs, although this metric reflects only the degree of linear relationship between predicted and behavioral FDMs (for additional discussion, see: Borji & Itti^5^; Bylinskii et al.^77^). Table S1 reports the results of an evaluation comparing model predictions of search FDMs to behavioral search FDMs using each of these metrics. The findings that we report in the main text in the context of scanpath prediction also hold in the case of FDM prediction. Specifically, the IRL-Hi-Low-C model outperformed the others, and did so for all three metrics. Additionally, the Detector-Hi model also performed relatively well in all the metrics, supporting our conclusion that a simple detector does a relatively good job in predicting fixations in visual search.

---

**Figure S1.**
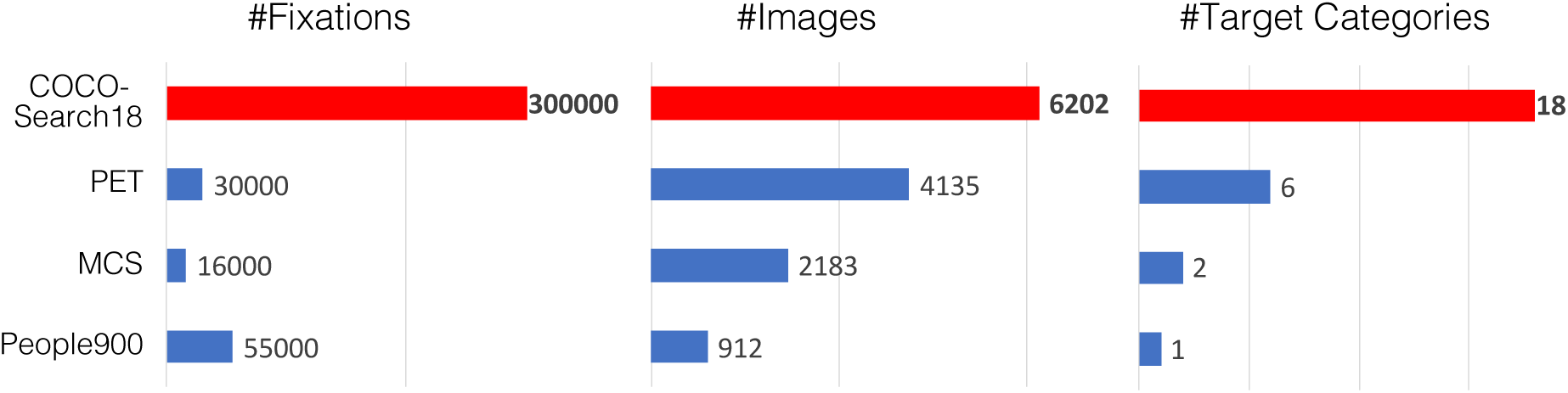
Comparisons between COCO-Search18 and other large-scale datasets of search behavior. COCO-Search18 is the largest in terms of number of fixations (*∼*300,000), number of target categories (18), and number of images (6,202).

**Figure S2.**
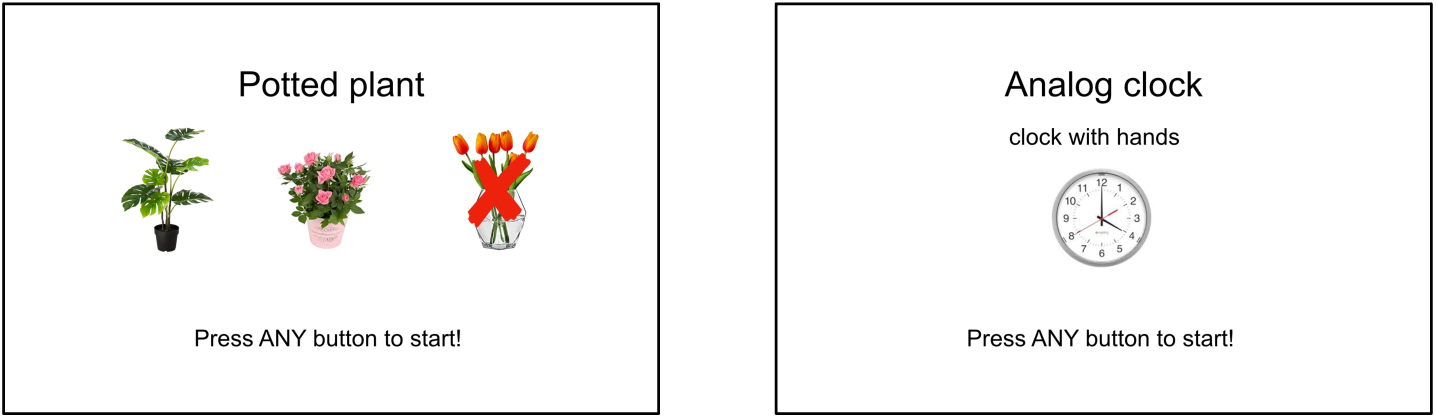
Examples of target-designation displays, shown for the potted-plant and analog clock targets, that preceded the block of trials for a given target category.

**Figure S3.**
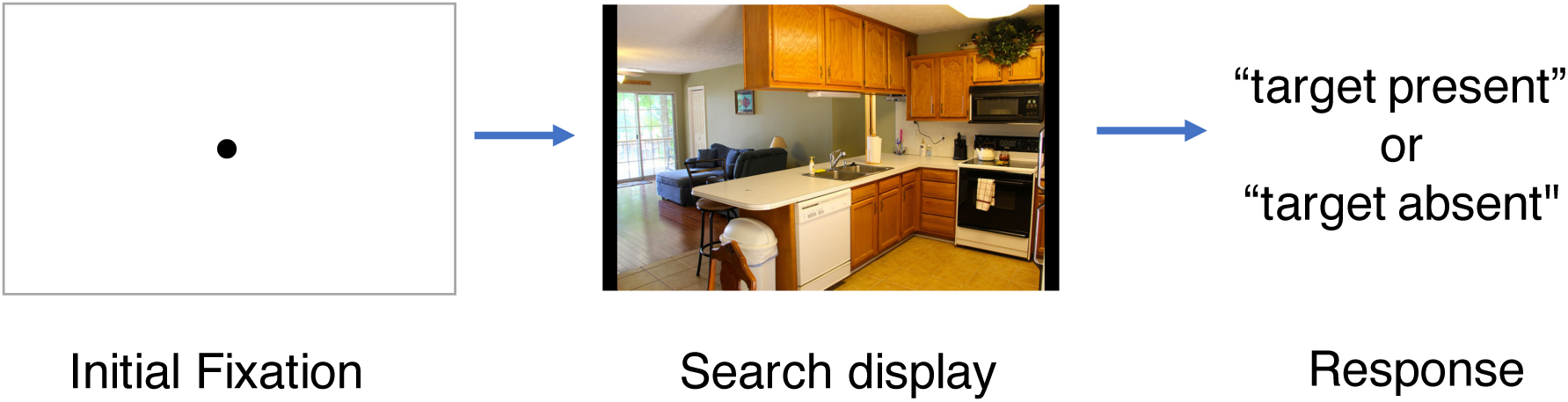
Example of the search procedure. Each trial began with a fixation dot appearing at the center of the screen. Participants would start a trial by pressing a button on a game-pad controller while carefully looking at the fixation dot. An image of a scene would then be displayed and the participant’s task was to make a speeded “yes” or “no” target-presence judgment by pressing the right or left triggers, respectively, of a game-pad controller.

**Figure S4.**
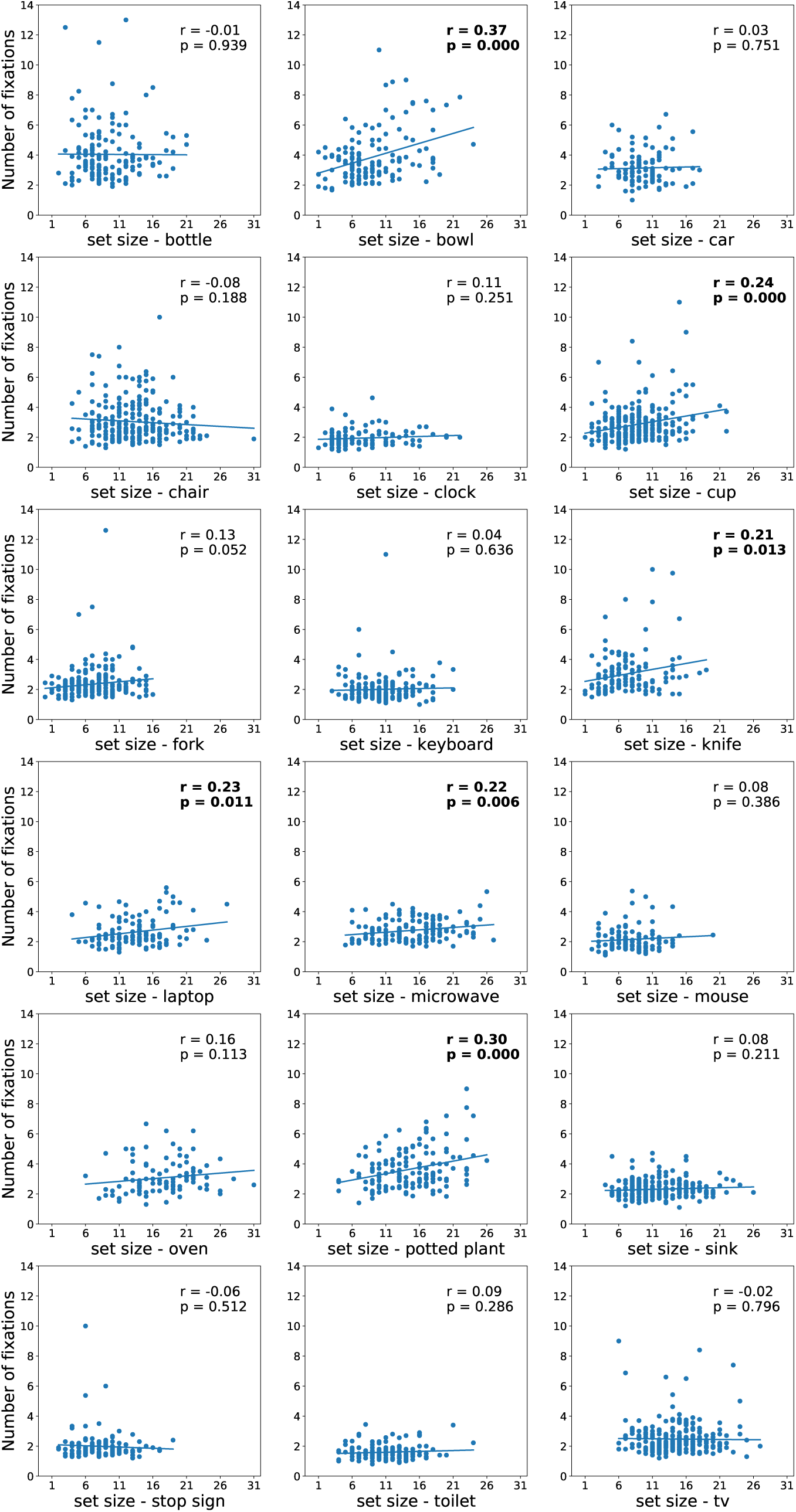
Number of fixations made on the target-present images plotted as a function of the set sizes of those images (using COCO object and stuff labels), averaged over participants and grouped by target category.

**Figure S5.**
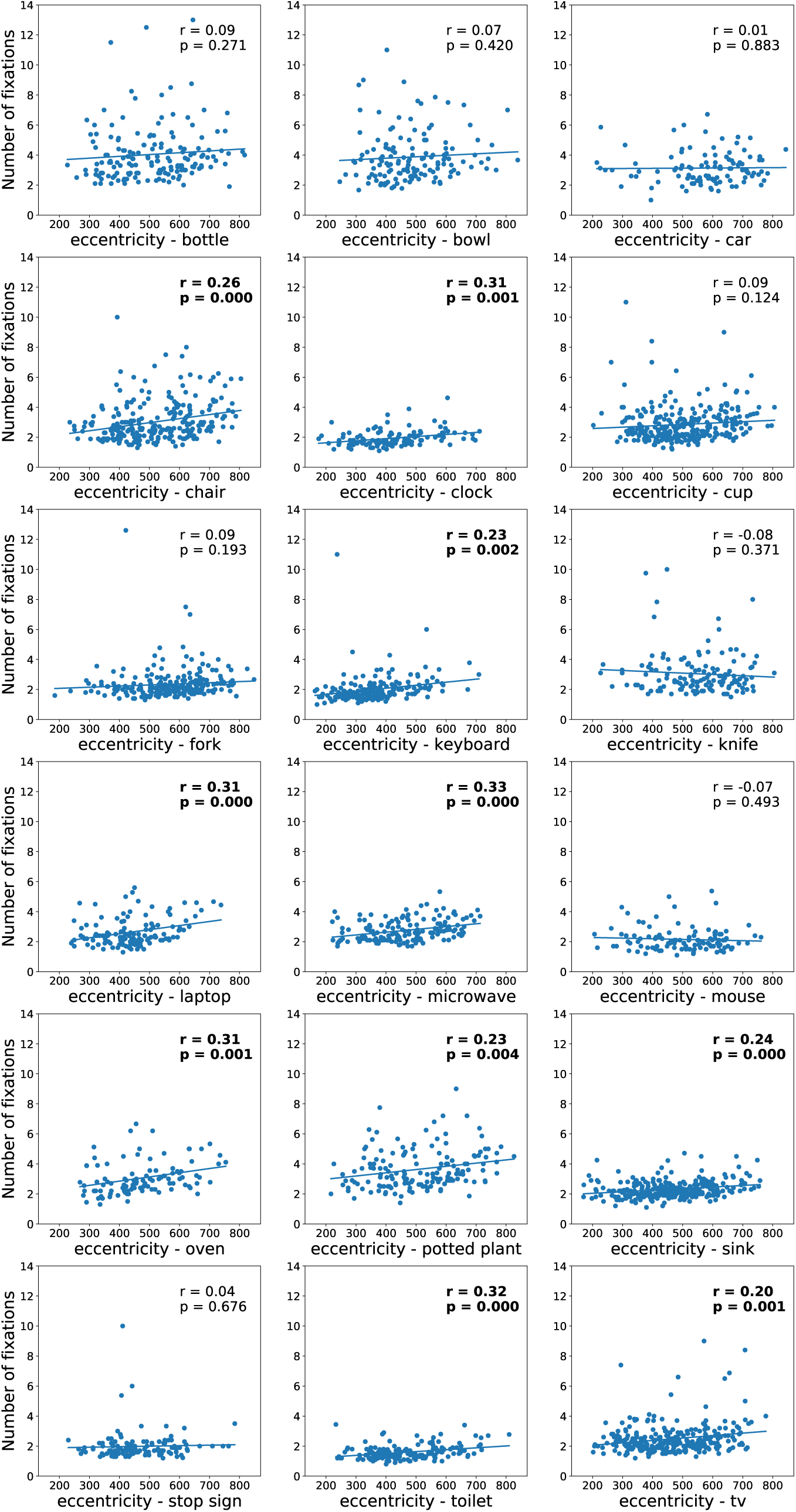
Number of fixations made on the target-present images plotted as a function of initial target eccentricity (using the center of the COCO bounding-box), averaged over participants and grouped by target category.

**Figure S6.**
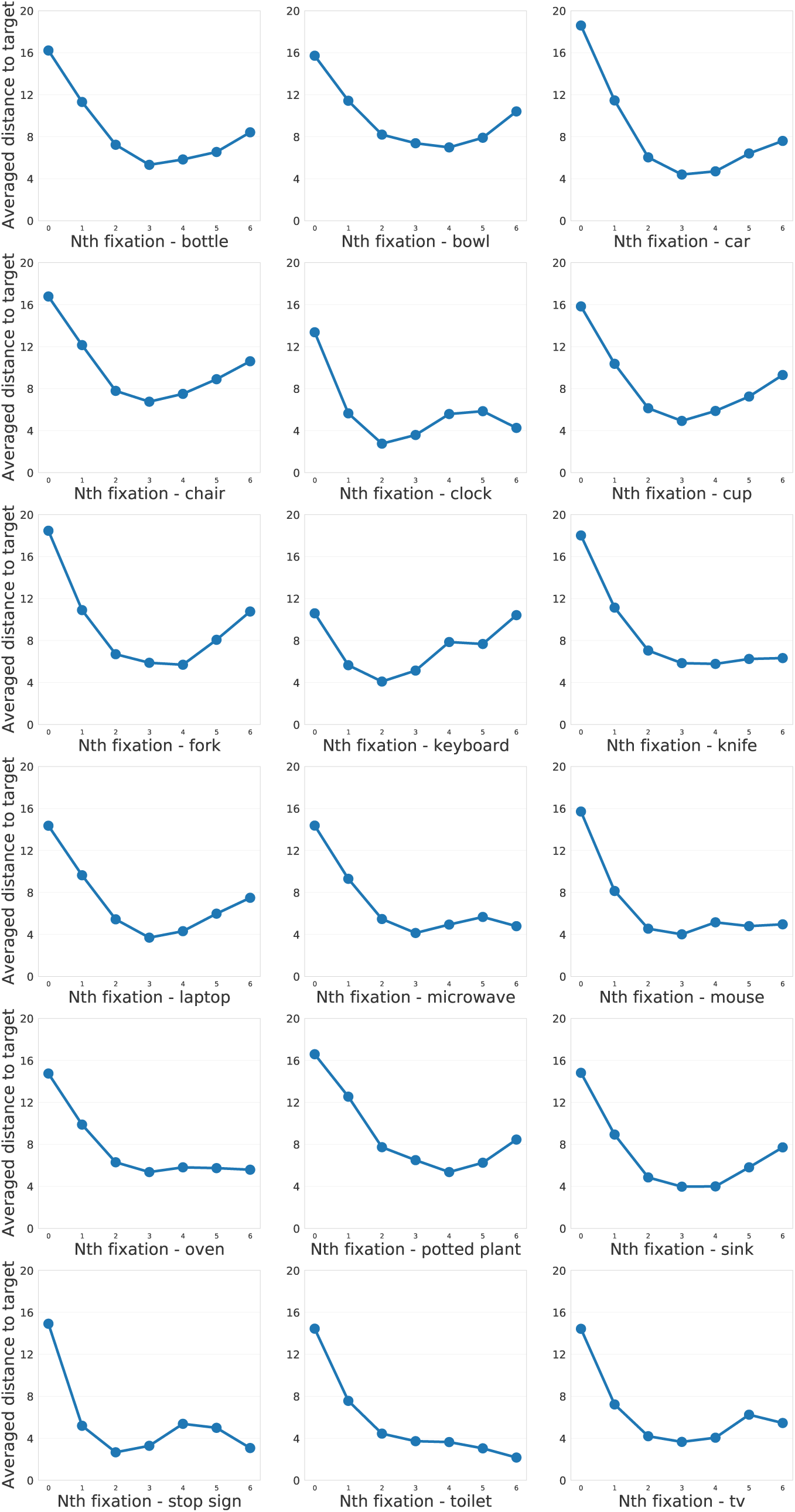
Averaged Euclidean distance (in visual angle) between gaze and the target’s center (using COCO bounding-box labels) over the first 6 saccades, grouped by target category.

**Figure S7.**
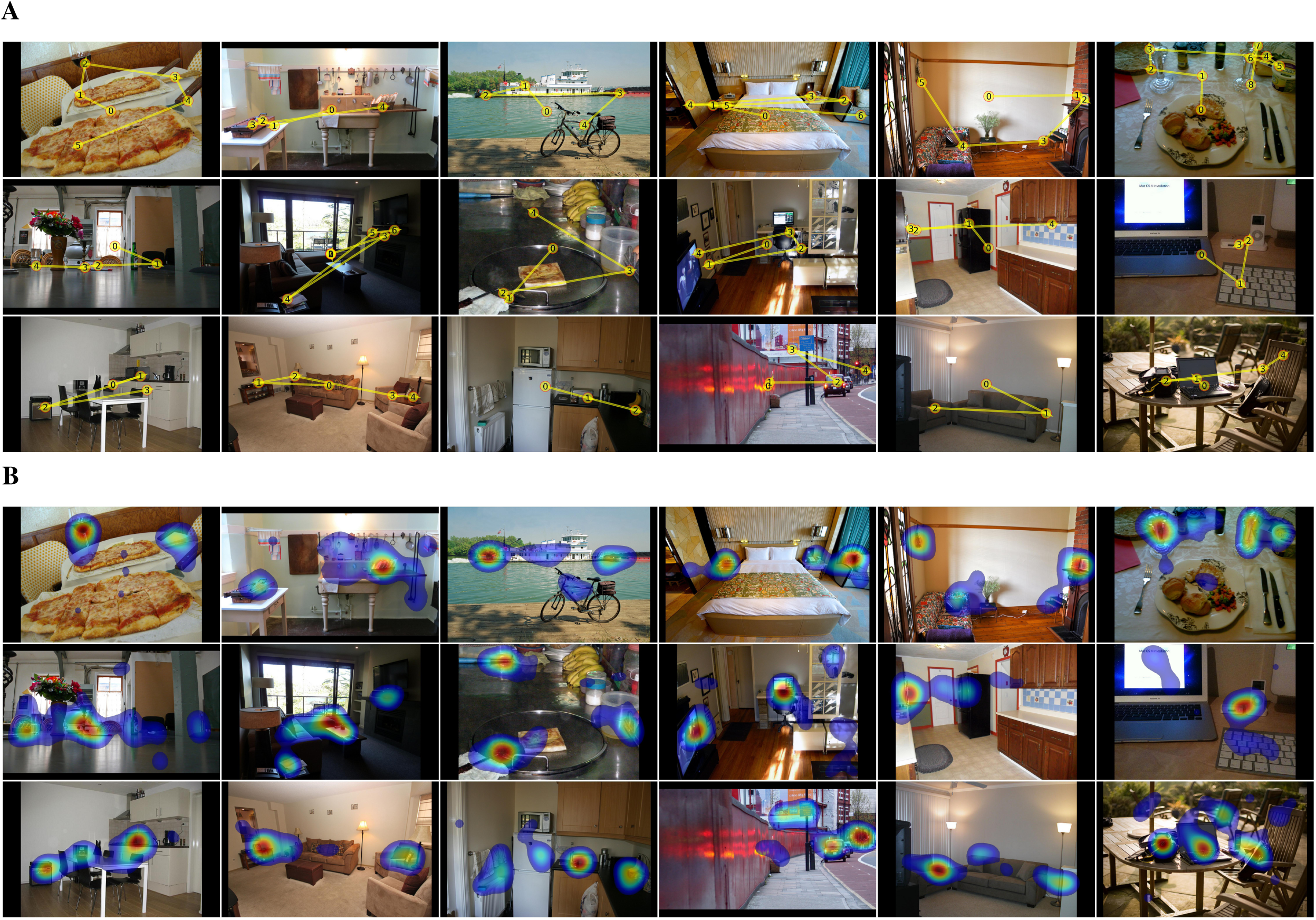
(A). Examples of a target-absent image for each of the 18 target categories. Yellow lines and numbered discs indicate a representative search scanpath from a single participant. From left to right, top to bottom: bottle, bowl, car, chair, (analog) clock, cup, fork, keyboard, knife, laptop, microwave, mouse, oven, potted plant, sink, stop sign, toilet, tv. (B). Examples of fixation density maps for the same target-absent images.

**Figure S8.**
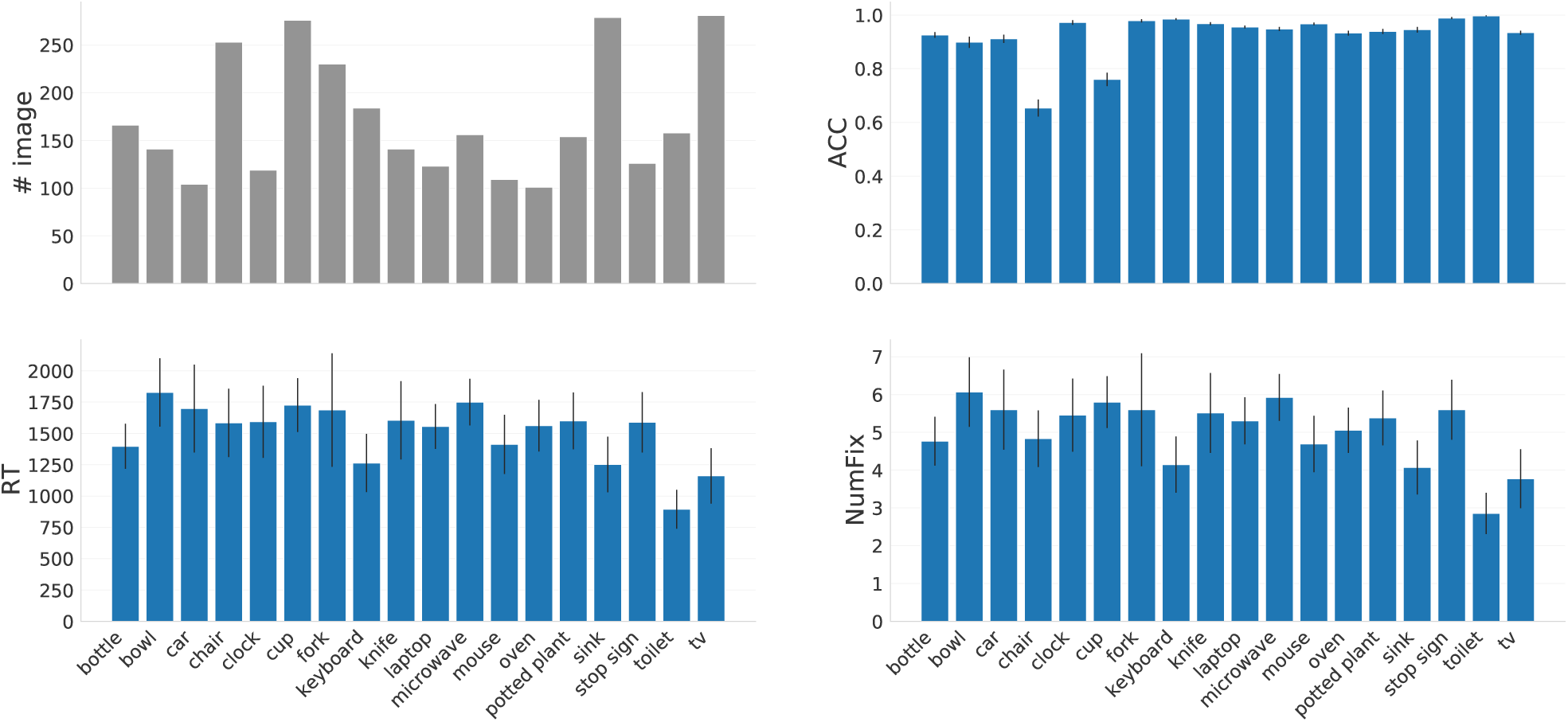
COCO-Search18 analyses for all 18 target categories in target-absent trials. Top: number of images in each category (gray), and response accuracy (ACC). Bottom: reaction time (RT) and number of fixations made before the button press (NumFix). Values are means over 10 participants, and error bars represent standard errors.

**Figure S9.**
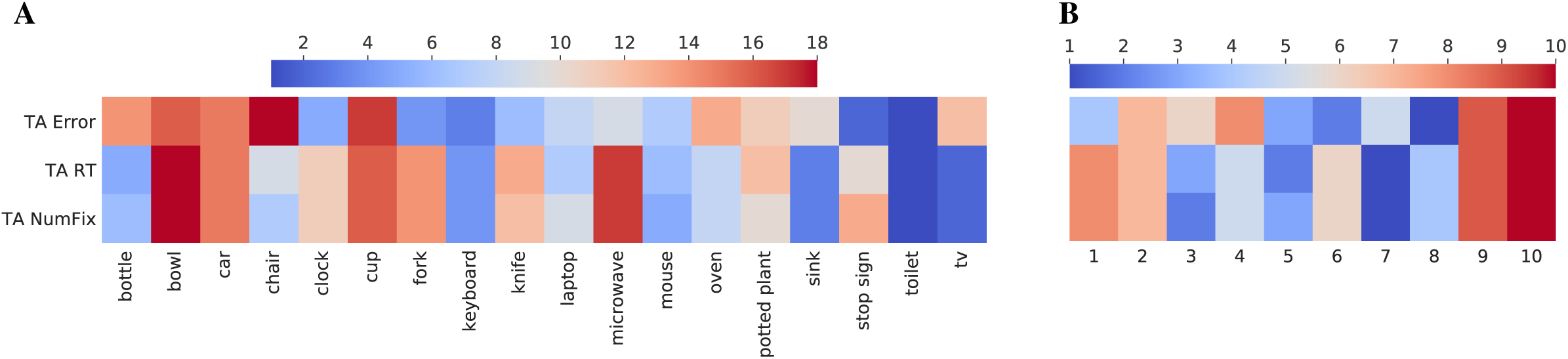
(A). Target-absent data, ranked [1-18] by target category (columns) and averaged over participants, shown for multiple performance measures (rows). These include: response error, reaction time (RT), and number of fixations (NumFix). Redder color indicates higher rank and harder search targets, bluer color indicates lower rank and easier search. (B) Target-absent data, now ranked by participant [1-10] and averaged over target category (columns). Performance measures and color coding are the same as in (A).

**Figure S10.**
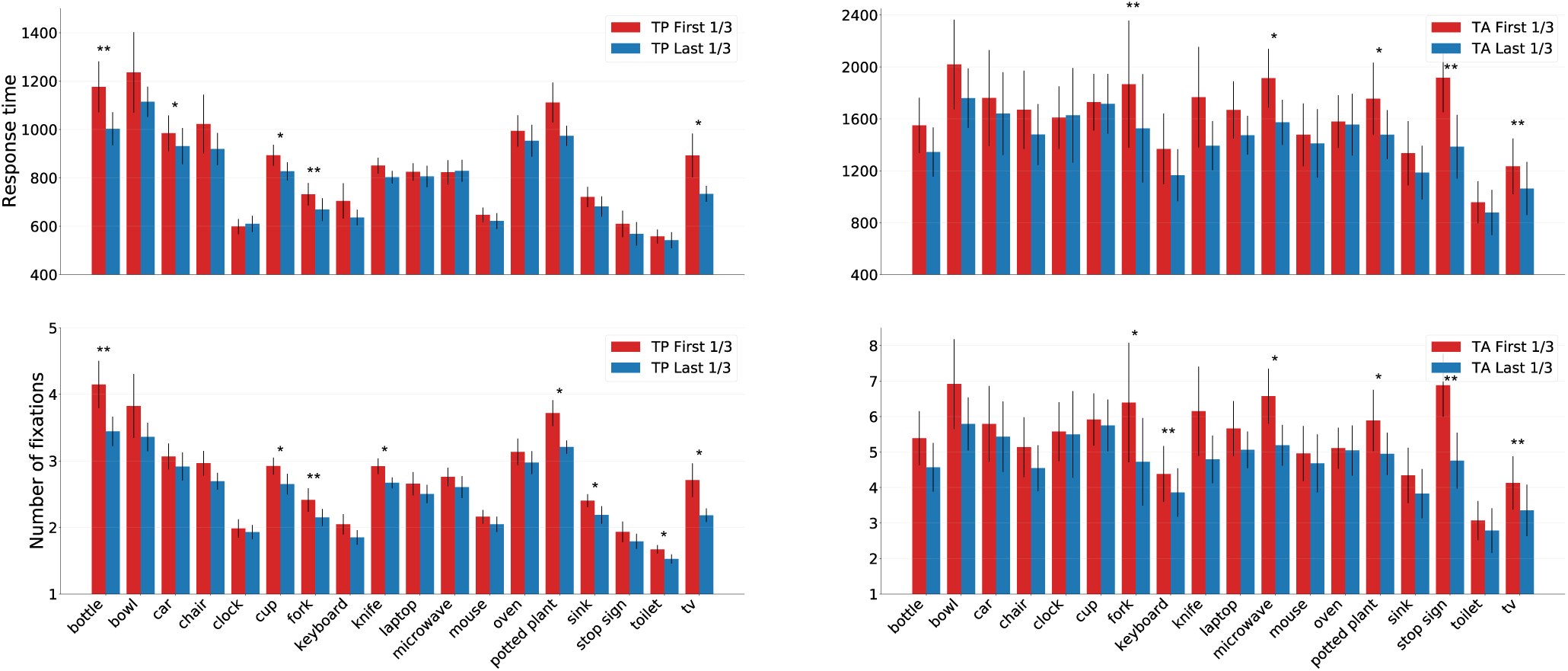
Practice effects, visualized as the difference in search performance between the red (first 1/3 of the trials) and the blue (last 1/3 of the trials) bars, grouped by the 18 target categories. The top row shows response time, and the bottom row shows the number of fixations before the button press. Target-present data are shown on the left, target-absent data are shown on the right. Only correct trials were included. *: *p* < .05, **: *p* < .01

**Figure S11.**
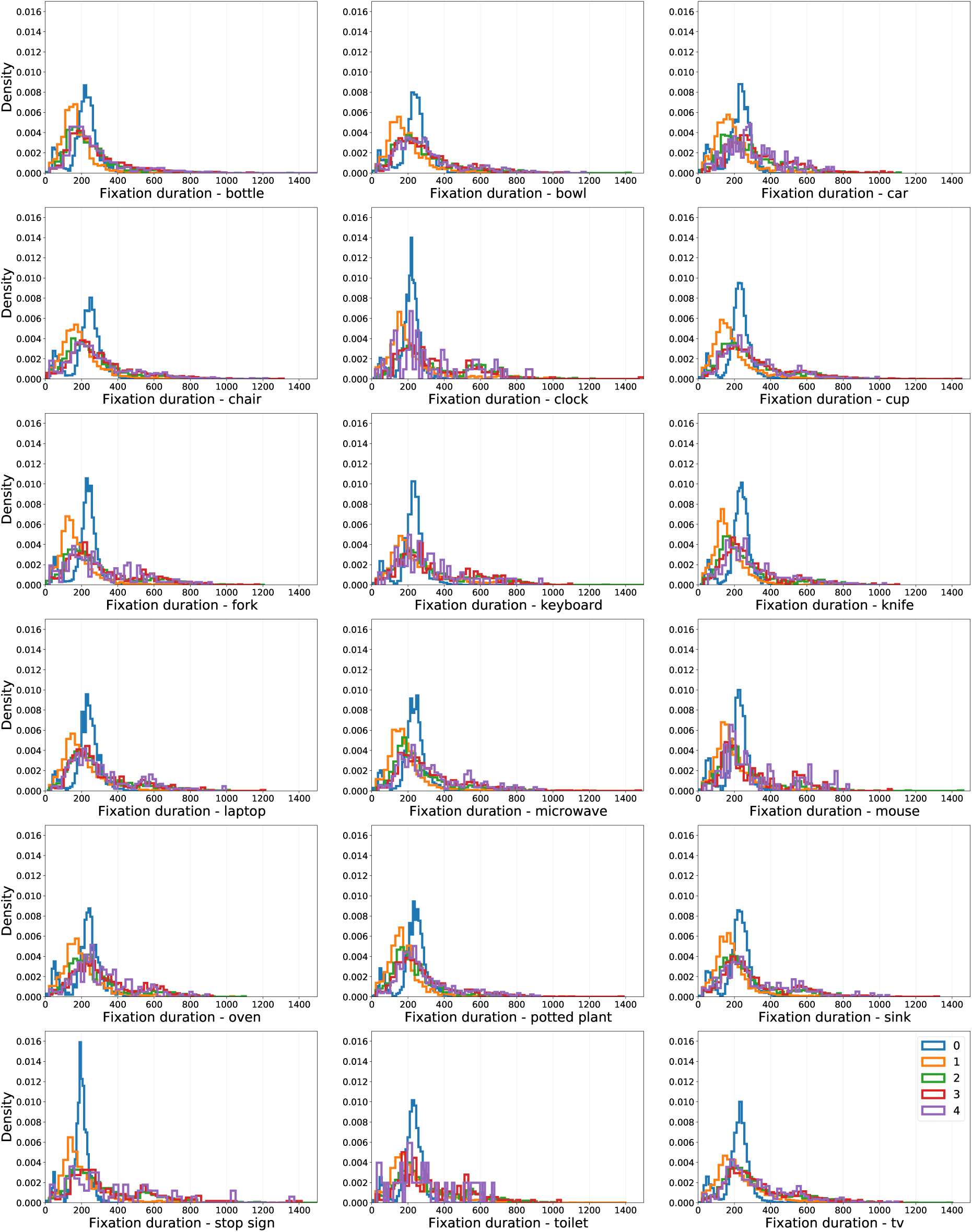
Density distributions of target-present fixation durations, plotted for each of the target categories (bin size = 50ms). The color lines refer to the initial fixation durations (0, blue), followed by the first four new fixations (1-4).

**Figure S12.**
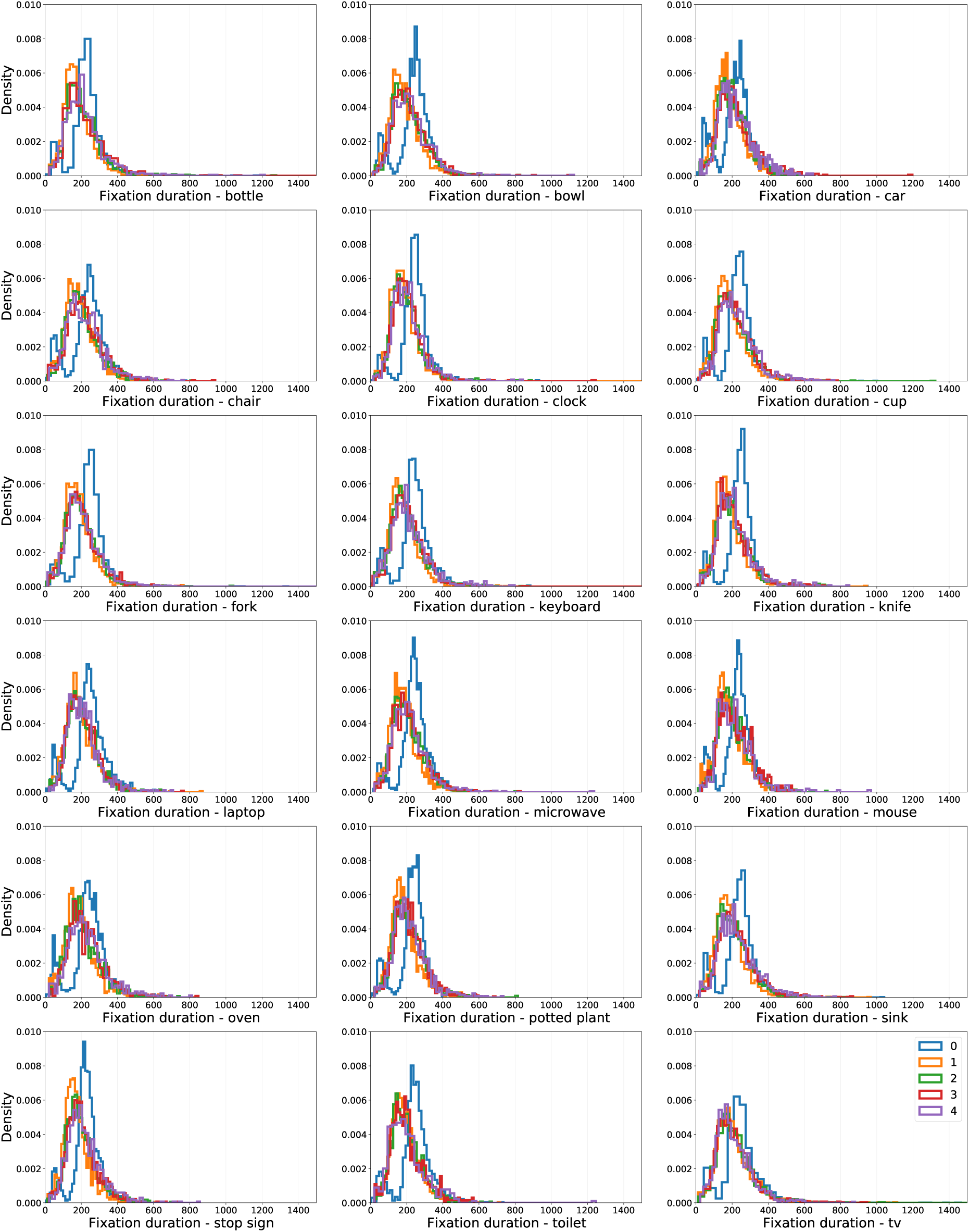
Density distributions of target-absent fixation durations, plotted for each of the target categories (bin size = 50ms). The color lines refer to the initial fixation durations (0, blue), followed by the first four new fixations (1-4).

**Figure S13.**
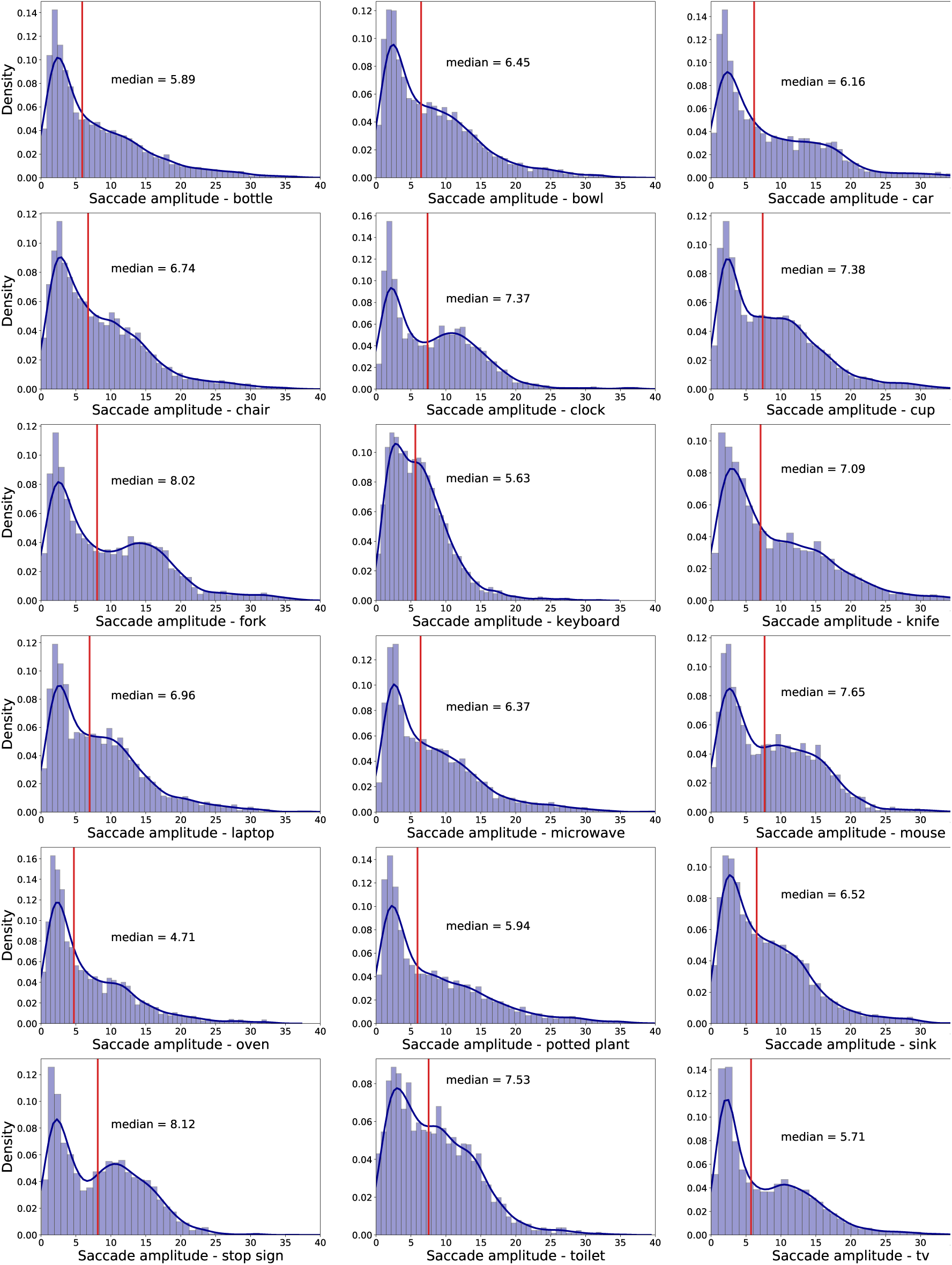
Density distributions of target-present saccade amplitudes (in visual angle), plotted by target category. Red vertical lines indicate median amplitudes. Dark blue lines represent Gaussian kernel density estimates.

**Figure S14.**
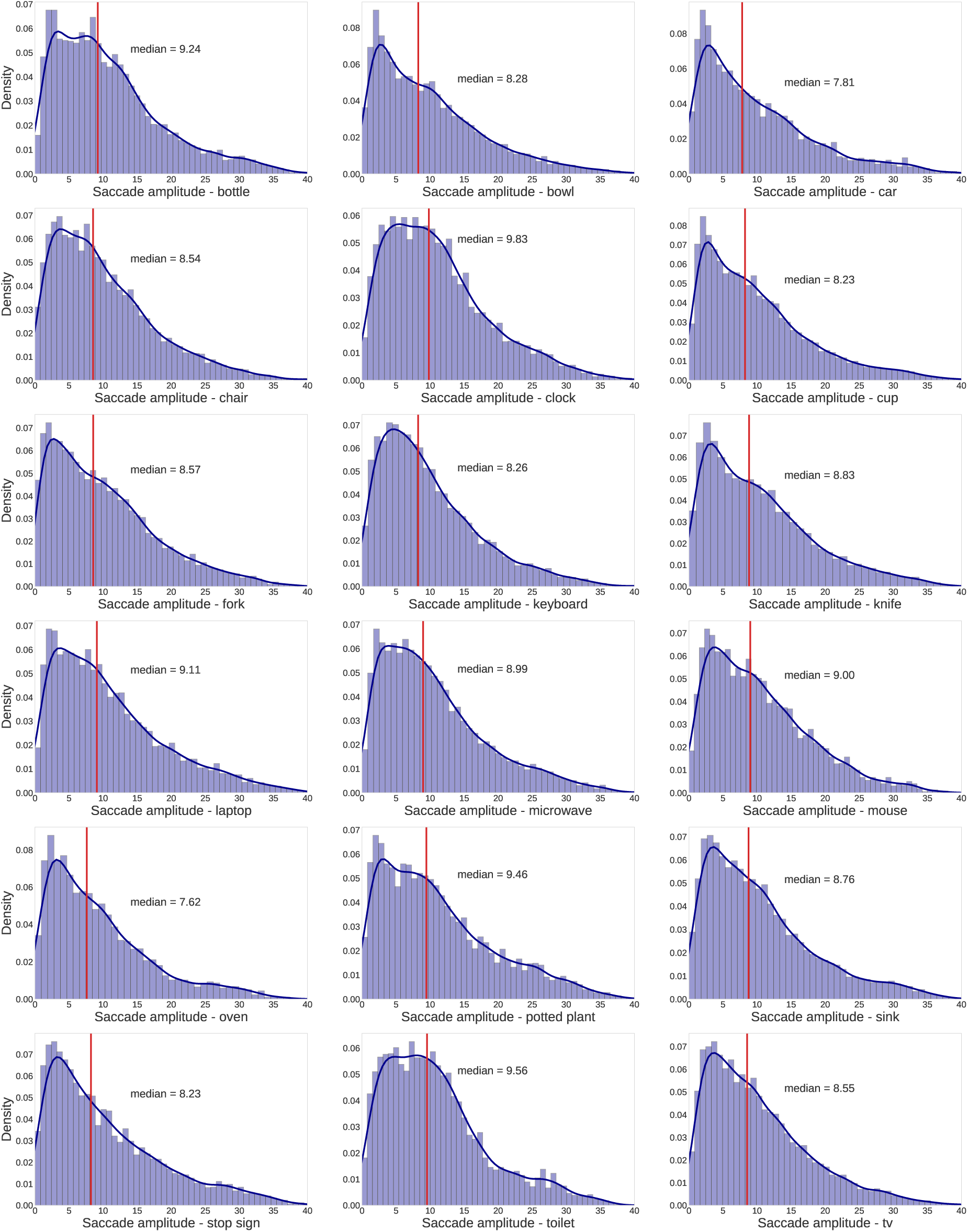
Density distributions of target-absent saccade amplitudes (in visual angle), plotted by target category. Red vertical lines indicate median amplitudes. Dark blue lines represent Gaussian kernel density estimates.

**Figure S15.**
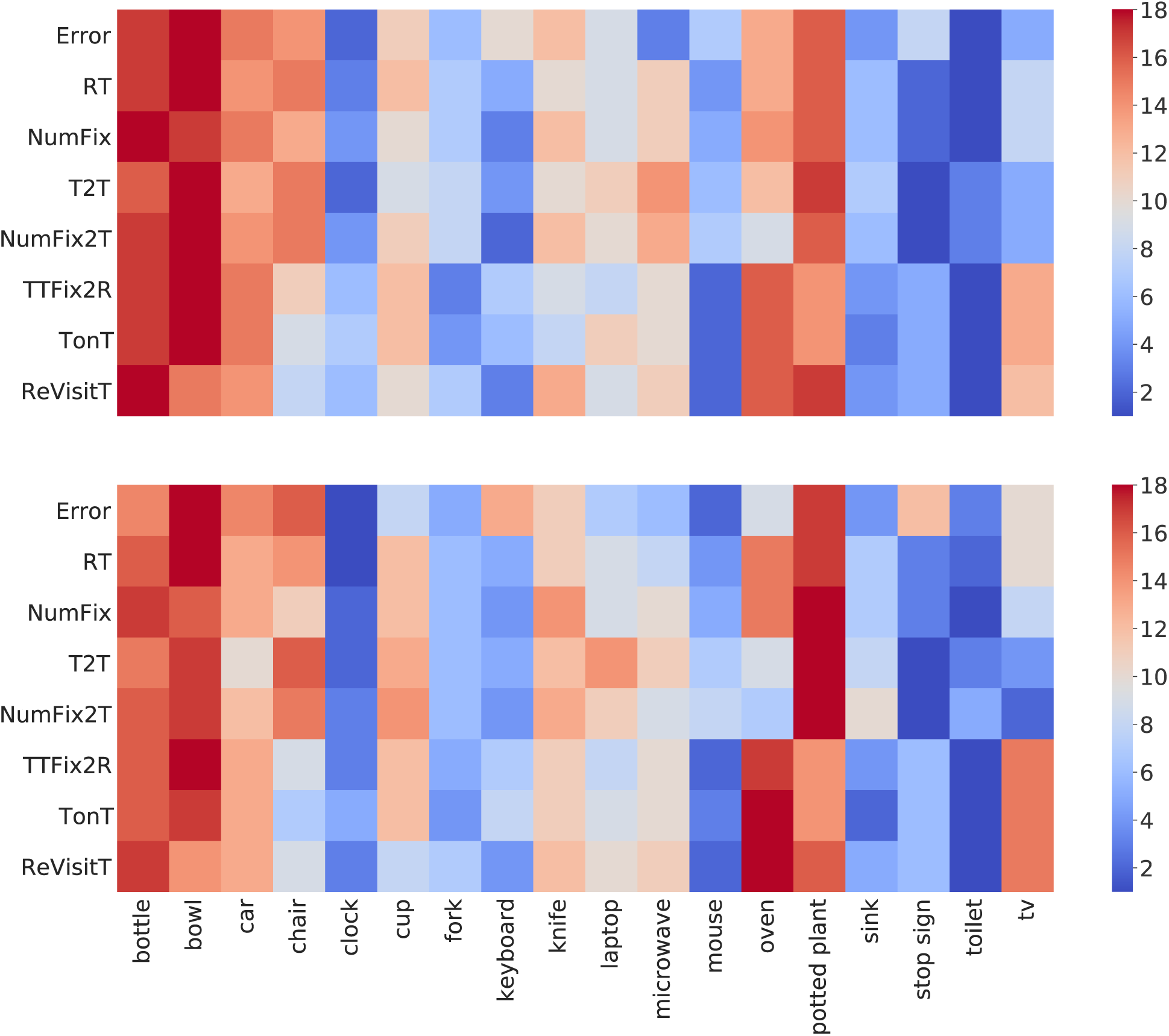
Target-present data, ranked by target category (1-18, columns) and shown for multiple performance measures (rows) in the trainval (top) and test (bottom) COCO-Search18 datasets. Redder color indicates higher rank and harder search targets, bluer color indicates lower rank and easier search. Measuers include: response error, reaction time (RT), number of fixations (NumFix), time to target (T2T), number of fixations to target (NumFix2T), time from first target fixation until response (TTFix2R), time spent fixating the target (TonT), and the number of target re-fixations (ReVisitT).

**Figure S16.**
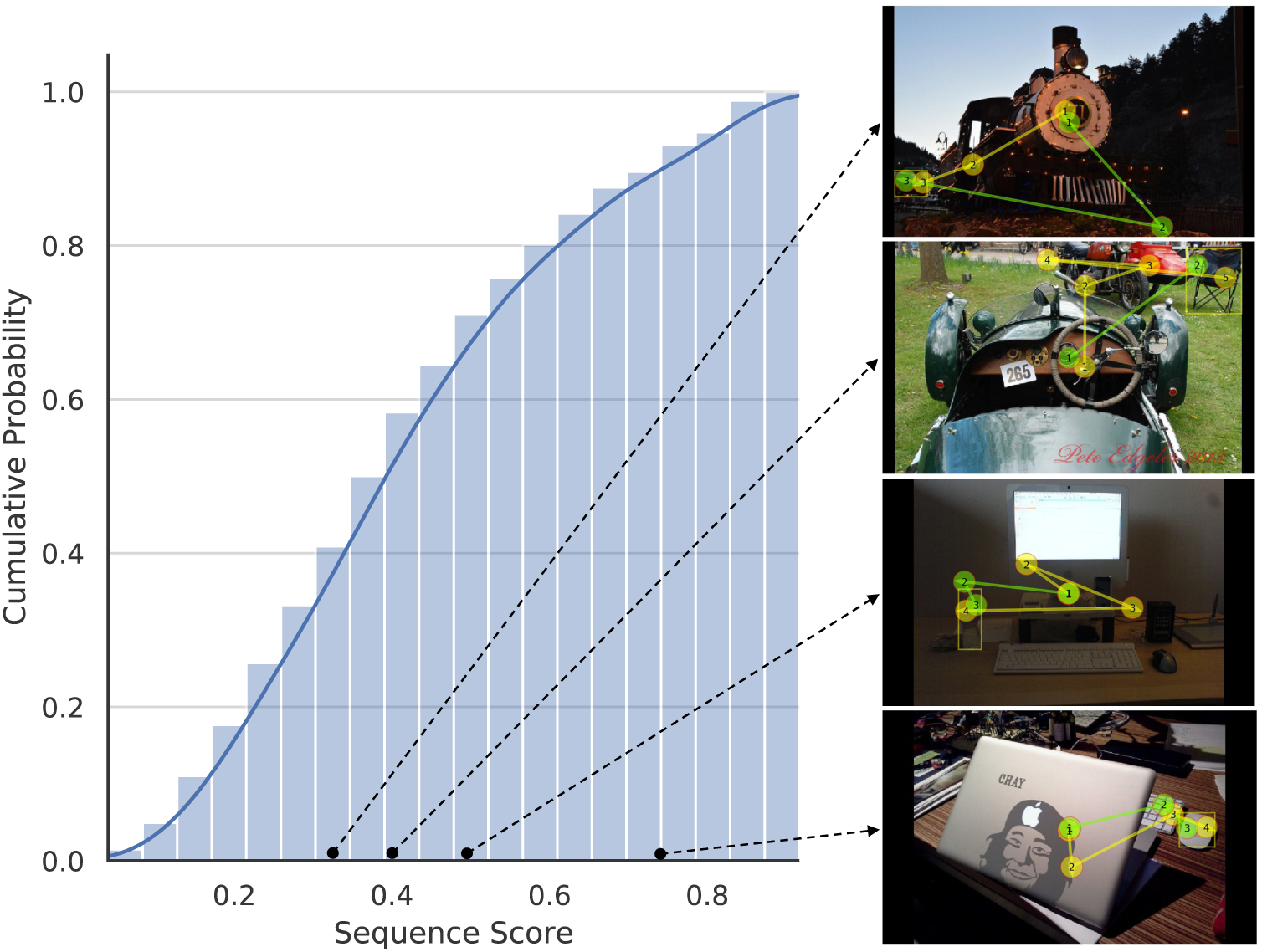
Left: cumulative distribution of average sequence scores computed between each scanpath generated by the IRL model and each behavioral scanpath for the test images of COCO-Search18. Right: Examples illustrating the scanpaths producing four different sequence scores. Behavioral scanpaths are colored in yellow, and the IRL-generated scanpaths are in green. Sequence scores for the four illustrated examples are 0.33, 0.40, 0.50, and 0.75, from top to bottom. Note that these results are from a slightly different version of the IRL model than the one reported here.

**Table S1.** Results from models (rows) predicting behavioral fixation-density maps (FDMs) using three spatial comparison metrics (columns), applied to the COCO-Search18 test images. “Human” refers to an oracle method whereby the FDM from half of the searchers was used to predict the FDM from the other half of the searchers. See the supplemental text for additional details about the spatial fixation comparison metrics.

|  | AUC ↑ | NSS ↑ | CC ↑ |
| --- | --- | --- | --- |
| Human | 0.675 | 3.396 | 0.356 |
| Random | 0.531 | 0.280 | 0.039 |
| Detector-Hi | 0.605 | 1.210 | 0.163 |
| Detector-Hi-Low | 0.575 | 0.792 | 0.105 |
| Deep Search-Hi | 0.620 | 1.122 | 0.153 |
| Deep Search-Hi-Low | 0.598 | 0.864 | 0.118 |
| IRL-ReT-C | 0.595 | 1.601 | 0.214 |
| IRL-Hi-Low-C | **0.628** | **1.806** | **0.246** |
| IRL-Hi-Low | 0.621 | 1.728 | 0.235 |

**Table S2.** *P* values from post-hoc t-tests (Bonferroni corrected) comparing predictive models (rows), averaged across the 18 target categories, for multiple scanpath metrics (columns). All *dfs* = 34. For decisively significant comparisons, the more predictive model is indicated in boldface.

| Compared Models | TFP-AUC | Probability Mismatch | Scanpath Ratio | Sequence Score | MultiMatch |  |  |  |
| --- | --- | --- | --- | --- | --- | --- | --- | --- |
|  |  |  |  |  | shape | direction | length | position |
| IRL-ReT-C vs. IRL-Hi-Low-C | *n.s.* | *n.s.* | *n.s.* | *n.s.* | *n.s.* | *n.s.* | *n.s.* | *n.s.* |
| IRL-ReT-C vs. IRL-Hi-Low | *n.s.* | *n.s.* | *n.s.* | *n.s.* | *n.s.* | *n.s.* | *n.s.* | *n.s.* |
| IRL-ReT-C vs. Detector-Hi | *n.s.* | *n.s.* | *n.s.* | *n.s.* | *n.s.* | *n.s.* | *n.s.* | *n.s.* |
| **IRL-ReT-C** vs. Detector-Hi-Low | .0017 | <.001 | <.001 | *n.s.* | .005 | .0686 | <.001 | .0039 |
| **IRL-ReT-C** vs. Deep Search-Hi | <.001 | <.001 | <.001 | *n.s.* | *n.s.* | <.001 | *n.s.* | *n.s.* |
| **IRL-ReT-C** vs. Deep Search-Hi-Low | <.001 | <.001 | <.001 | .0587 | *n.s.* | <.001 | *n.s.* | *n.s.* |
| IRL-Hi-Low-C vs. IRL-Hi-Low | *n.s.* | *n.s.* | *n.s.* | *n.s.* | *n.s.* | *n.s.* | *n.s.* | *n.s.* |
| IRL-Hi-Low-C vs. Detector-Hi | *n.s.* | *n.s.* | .0653 | *n.s.* | *n.s.* | *n.s.* | .0235 | *n.s.* |
| **IRL-Hi-Low-C** vs. Detector-Hi-Low | <.001 | <.001 | <.001 | *n.s.* | <.001 | .0515 | <.001 | <.001 |
| **IRL-Hi-Low-C** vs. Deep Search-Hi | <.001 | <.001 | <.001 | *n.s.* | *n.s.* | <.001 | *n.s.* | *n.s.* |
| **IRL-Hi-Low-C** vs. Deep Search-Hi-Low | <.001 | <.001 | <.001 | .0559 | .0298 | <.001 | *n.s.* | .0110 |
| IRL-Hi-Low vs. Detector-Hi | *n.s.* | *n.s.* | .0151 | *n.s.* | *n.s.* | *n.s.* | .0206 | *n.s.* |
| **IRL-Hi-Low** vs. Detector-Hi-Low | <.001 | <.001 | <.001 | *n.s.* | <.001 | .0539 | <.001 | <.001 |
| **IRL-Hi-Low** vs. Deep Search-Hi | <.001 | <.001 | <.001 | *n.s.* | *n.s.* | <.001 | *n.s.* | *n.s.* |
| **IRL-Hi-Low** vs. Deep Search-Hi-Low | <.001 | <.001 | <.001 | .0506 | *n.s.* | <.001 | *n.s.* | .0029 |
| **Detector-Hi** vs. Detector-Hi-Low | .0019 | <.001 | .0086 | *n.s.* | *n.s.* | *n.s.* | *n.s.* | .0150 |
| **Detector-Hi** vs. Deep Search-Hi | <.001 | <.001 | <.001 | *n.s.* | *n.s.* | .0013 | <.001 | *n.s.* |
| **Detector-Hi** vs. Deep Search-Hi-Low | <.001 | <.001 | <.001 | .0755 | *n.s.* | <.001 | <.001 | *n.s.* |
| Detector-Hi-Low vs. Deep Search-Hi | *n.s.* | *n.s.* | *n.s.* | *n.s.* | <.001 | *n.s.* | <.001 | <.001 |
| Detector-Hi-Low vs. Deep Search-Hi-Low | *n.s.* | .0275 | *n.s.* | *n.s.* | .0446 | *n.s.* | <.001 | .0511 |
| Deep Search-Hi vs. Deep Search-Hi-Low | *n.s.* | *n.s.* | *n.s.* | *n.s.* | *n.s.* | *n.s.* | *n.s.* | .0778 |

